# Loss of cardiac PFKFB2 drives Metabolic, Functional, and Electrophysiological Remodeling in the Heart

**DOI:** 10.1101/2023.11.22.568379

**Authors:** Kylene M. Harold, Satoshi Matsuzaki, Atul Pranay, Brooke L. Loveland, Albert Batushansky, Maria F. Mendez Garcia, Craig Eyster, Stavros Stavrakis, Ying Ann Chiao, Michael Kinter, Kenneth M. Humphries

**Affiliations:** Aging and Metabolism Research Program, Oklahoma Medical Research Foundation, Oklahoma City, OK; Department of Biochemistry and Molecular Physiology, University of Oklahoma Health Sciences Center, Oklahoma City, OK; Ilse Katz Institute for Nanoscale Science & Technology, Ben-Gurion University of the Negev, Beer Sheva, Israel; Department of Medicine, Section of Cardiovascular Medicine, University of Oklahoma Health Sciences Center, Oklahoma City, Oklahoma, USA

**Keywords:** Glycolysis, glucose, metabolism, echocardiography, electrocardiography

## Abstract

**Background:** Phosphofructo-2-kinase/fructose-2,6-bisphosphatase (PFK-2) is a critical glycolytic regulator responsible for upregulation of glycolysis in response to insulin and adrenergic signaling. PFKFB2, the cardiac isoform of PFK-2, is degraded in the heart in the absence of insulin signaling, contributing to diabetes-induced cardiac metabolic inflexibility. However, previous studies have not examined how the loss of PFKFB2 affects global cardiac metabolism and function.

**Methods:** To address this, we have generated a mouse model with a cardiomyocyte-specific knockout of PFKFB2 (cKO). Using 9-month-old cKO and control (CON) mice, we characterized impacts of PFKFB2 on cardiac metabolism, function, and electrophysiology.

**Results:** cKO mice have a shortened lifespan of 9 months. Metabolically, cKO mice are characterized by increased glycolytic enzyme abundance and pyruvate dehydrogenase (PDH) activity, as well as decreased mitochondrial abundance and beta oxidation, suggesting a shift toward glucose metabolism. This was supported by a decrease in the ratio of palmitoyl carnitine to pyruvate-dependent mitochondrial respiration in cKO relative to CON animals. Metabolomic, proteomic, and western blot data support the activation of ancillary glucose metabolism, including pentose phosphate and hexosamine biosynthesis pathways. Physiologically, cKO animals exhibited impaired systolic function and left ventricular (LV) dilation, represented by reduced fractional shortening and increased LV internal diameter, respectively. This was accompanied by electrophysiological alterations including increased QT interval and other metrics of delayed ventricular conduction.

**Conclusions:** Loss of PFKFB2 results in metabolic remodeling marked by cardiac ancillary pathway activation. This could delineate an underpinning of pathologic changes to mechanical and electrical function in the heart.

**Clinical Perspective:** *What is New?:*

- We have generated a novel cardiomyocyte-specific knockout model of PFKFB2, the cardiac isoform of the primary glycolytic regulator Phosphofructokinase-2 (cKO).
- The cKO model demonstrates that loss of cardiac PFKFB2 drives metabolic reprogramming and shunting of glucose metabolites to ancillary metabolic pathways.
- The loss of cardiac PFKFB2 promotes electrophysiological and functional remodeling in the cKO heart.

*What are the Clinical Implications?:*

- PFKFB2 is degraded in the absence of insulin signaling, making its loss particularly relevant to diabetes and the pathophysiology of diabetic cardiomyopathy.
- Changes which we observe in the cKO model are consistent with those often observed in diabetes and heart failure of other etiologies.
- Defining PFKFB2 loss as a driver of cardiac pathogenesis identifies it as a target for future investigation and potential therapeutic intervention.

## Introduction

Due to its critical role in maintaining perfusion, the heart must sustain consistent energetic homeostasis. This requires sufficient adaptive means to upregulate energy production under stress conditions when requirements increase. Further, the heart must be capable of acutely remodeling its metabolic phenotype when substrate availability is altered. One mechanism through which it achieves these outcomes relies on phosphofructo-2-kinase/fructose 2,6-bisphosphatase (PFK-2), a glycolytic regulator with an imperative role in modulating phosphofructokinase 1 (PFK-1), the first rate-limiting enzyme of glycolysis [1, 2]. PFK-2 is a bifunctional enzyme, capable of both phosphorylating fructose 6-phosphate to produce fructose 2,6-bisphosphate or catalyzing the reverse reaction [3]. When the cardiac isoform of PFK-2 (PFKFB2) is phosphorylated by adrenergic or insulin signaling pathways, the kinase form dominates, catalyzing the forward reaction and driving increased abundance of fructose 2,6-bisphosphate, the strongest allosteric activator of PFK-1 [4, 5]. This allows for an increase in glycolysis in response to stress or nutrient sensing. Reciprocally, when dephosphorylated, PFK-2 has dominant phosphatase activity which decreases fructose 2,6-bisphosphate levels and PFK-1 activation.

There are several isoforms of PFK-2, with the PFKFB2 isoform predominantly expressed in the heart [6]. We have previously shown that abundance of PFKFB2, referred to in our previous reports as PFK-2, is dynamically regulated by insulin signaling [2, 7]. PFKFB2 content is sustained in response to insulin signaling but is rapidly degraded by both proteasomal and autophagic mechanisms in the absence of insulin signaling [2]. Thus, in a murine streptozotocin-induced type 1 diabetic model, cardiac PFKFB2 content is dramatically decreased and remaining enzyme is unable to be phosphorylated by adrenergic cues [2, 8]. This maladaptation contributes to the metabolic inflexibility of the diabetic heart. However, it is unknown if or how the loss of PFKFB2 directly affects intermediate metabolism and cardiac function. To address this question, we developed a mouse model with a cardiomyocyte-specific knockout of PFKFB2 (cKO).

Diabetes has been linked to atrial and ventricular arrhythmias [9, 10]. Established contributions of cardiac autonomic neuropathy and metabolic stress-driven remodeling only partially explain this metabolic etiology and why changes to parameters such as glycemic status are closely associated with arrhythmogenic propensity. A coupling of glucose oxidation with intracellular ion handling in the cardiomyocyte has been noted [11]. This is likely due in part to activation of ancillary pathways such as the hexosamine biosynthesis pathway (HBP) and other processes which directly or indirectly alter intracellular ion handling [12-14]. However, the underlying mechanisms driving these changes in metabolic disease remain to be fully elucidated. PFK-2 mediation of glycolytic regulation could provide insight into these areas. However, the impacts of alterations at the PFKFB2/PFK-1 regulatory nexus on cardiac electrophysiology have not been previously explored. The relationship between heart failure and arrhythmia is complex, and shared mechanisms such as HBP and CAMKII activation underlie electrophysiological and mechanical dysfunction in diseases of metabolic etiology [12, 15-17]. While overexpression of PFK-2 in type 2 diabetes protects against functional changes [18], mechanisms by which PFK-2 abundance and activity may contribute to proper cardiac function are not completely understood.

In this study, we sought to determine how the loss of PFKFB2 affects cardiac function and metabolism. We demonstrate metabolic remodeling that diminishes mitochondrial abundance but increases glucose oxidation. There is also an increase in alternate glucose metabolism through ancillary pathways and an increase in nutrient signaling enzymes Akt and AMPK. These metabolic adaptations are associated with decreased cardiac function and electrophysiological changes. The pathological alterations are consistent with those observed in a subset of patients with diabetes, highlighting the importance of proper metabolic regulation at the PFKFB2/PFK-1 nexus [19, 20].

## Methods

### Animals

All animal procedures were approved by the Oklahoma Medical Research Foundation Animal Care and Use Committee. Animals were maintained on a standard *ad libitum* lab chow diet. Mice on a C57BL/6J background were engineered (ViewSolid Biotech, Oklahoma City) to have two flox sites such that exon 2 of *Pfkfb2* (NCBI gene ID: 18640; Chromosome 1: 130,689,182-130,729,253 reverse strand) is excised upon Cre recombination. Exon 2 contains the transcriptional start site. Homozygous *Pfkfb2^flox/flox^* female mice without Cre were bred to *Pfkfb2^flox/flox^* mice heterozygous for Cre expressed under the myosin heavy chain 6 promoter (cKO) (The Jackson Laboratory, Strain 011038). This produced cKO and litter-matched control (CON) mice at a 1:1 (mendelian) ratio, suggesting little to no embryonic lethality. Due to a sudden cardiac death phenotype observed early in the characterization of this model, occurring at an average age of 274 ± 67 days (n=30), a 9-month terminal time point was chosen for all aspects of this study. Animals were euthanized via cervical dislocation or cardiac excision under isoflurane-induced general anesthesia following a 3 hour fast. Due to an absence of notable differences between sexes in preliminary experiments, a combination of male and female mice was used throughout the study.

### Homogenate Preparation and Mitochondrial Isolation

Animals were euthanized by cervical dislocation and hearts were perfused with mitochondrial isolation buffer (210 mM Mannitol, 70 mM Sucrose, 5 mM MOPS, 1 mM EDTA; pH 7.4) prior to excision. Hearts were homogenized (Transi-Stir-Q-2826, Talboys Engineering Corp.) in 5 mL of mitochondrial isolation buffer and centrifuged at 550 x g for 5 minutes at 4°C, after which 200 µL of supernatant was collected as homogenate. Remaining supernatant was centrifuged at 10000 x g for 10 minutes at 4°C. The resulting mitochondrial pellet was resuspended in 60 µL mitochondrial isolation buffer. BCA was used to quantify protein concentrations of both the mitochondrial suspension and homogenate.

### Mitochondrial Subfractionation

The mitochondrial subfractionation procedure was derived from a previous literature describing this technique [21, 22] and optimized for cardiac mitochondria. Following mitochondrial isolation and protein determination, mitochondria were diluted to approximately 1 mg/mL in TD buffer (50.0 mM Trizma base, 274.1 mM NaCl, 20.1 mM KCl, 14.0 mM Na_2_HPO_4_). Fifty microliter aliquots of the diluted mitochondria from each sample were incubated for 1h at room temperature in the following treatment groups: untreated, Proteinase K (≥0.075 units/mL), Proteinase K (≥0.15 units/mL), Proteinase K (≥0.075 units/mL) + 1.0% Triton X-100, or Proteinase K (≥0.15 units/mL) + 1.0% Triton X-100. Following incubation, 2 mM phenylmethylsulfonyl fluoride was added and samples were suspended in lithium dodecyl sulfate (LDS) sample buffer to prepare for western blot.

### Mitochondrial Respiration

Following isolation and suspension, mitochondria were further diluted to 0.25 mg/mL in OXPHOS buffer (210 mM mannitol, 70 mM sucrose, 10 mM MOPS, and 5 mM K_2_HPO_4_; pH 7.4), with 0.5 mg/mL BSA. Respiration (State 2) was initiated with either 0.1 mM pyruvate and 1.0 mM malate, or 30 µM palmitoyl carnitine and 1.0 mM malate. A fluorescence lifetime-based dissolved oxygen monitor system (Instech) was utilized to quantify oxygen consumption. Following 2 minutes of State 2 measurement, 0.5 mM ADP was added to induce State 3 respiration.

### Pyruvate Dehydrogenase Activity Assay

Mitochondria were isolated from cardiac tissue and suspended in 60 µL mitochondrial isolation buffer as described herein. The mitochondrial suspension was further diluted 3:100 in OXPHOS buffer (see above) with or without 0.25 mM pyruvate and 2.5 mM malate for activated or inactivated assays respectively. Following a 2 minute incubation, this solution was further diluted 1:5 in 25 mM MOPS with 0.05% Triton X-100. One mM NAD^+^, 2.5 mM pyruvate, 200 µM thiamine pyrophosphate, 100 µM CoASH, and 5.0 mM MgCl_2_ were then added and NAD^+^ reduction was measured as a proxy for PDH activity. NAD^+^ reduction was quantified spectrophotometrically as NADH absorbance at 340 nm every 5 seconds for 275 seconds following a 25 second incubation. Activities were normalized to protein concentrations of the original mitochondrial suspension, obtained by BCA.

### Western Blot

Western blot analysis was performed for whole heart tissue homogenates and isolated mitochondria as previously described [23]. Briefly, samples were prepared using LDS sample buffer (Thermo Fisher Scientific) with addition of a protease and phosphatase inhibitor cocktail (Halt^TM^, Thermo Fisher Scientific), after which they were run on 4-12% Bis-Tris gels (Thermo Fisher Scientific) at 200 V for 45 minutes. Protein was then transferred to nitrocellulose membranes at 30 V for 1 hour. Total protein was stained (Revert Total Protein Stain, LI-COR) and imaged (Odyssey CLx, LI-COR) for later normalization. Blots were blocked for 45 minutes (Intercept Blocking Buffer, LI-COR) and incubated in primary antibody for 1 hour at room temperature or overnight at 4°C. Primary antibodies and their concentrations used are depicted in **Table 1**. Signal from corresponding anti-rabbit (IRDye 800CW, LI-COR) or anti-mouse (IRDye 680RD, LI-COR) secondary antibodies was imaged (Odyssey CLx, LI-COR) and quantified via Image Studio (Version 5.2, LI-COR).

**Table 1.**
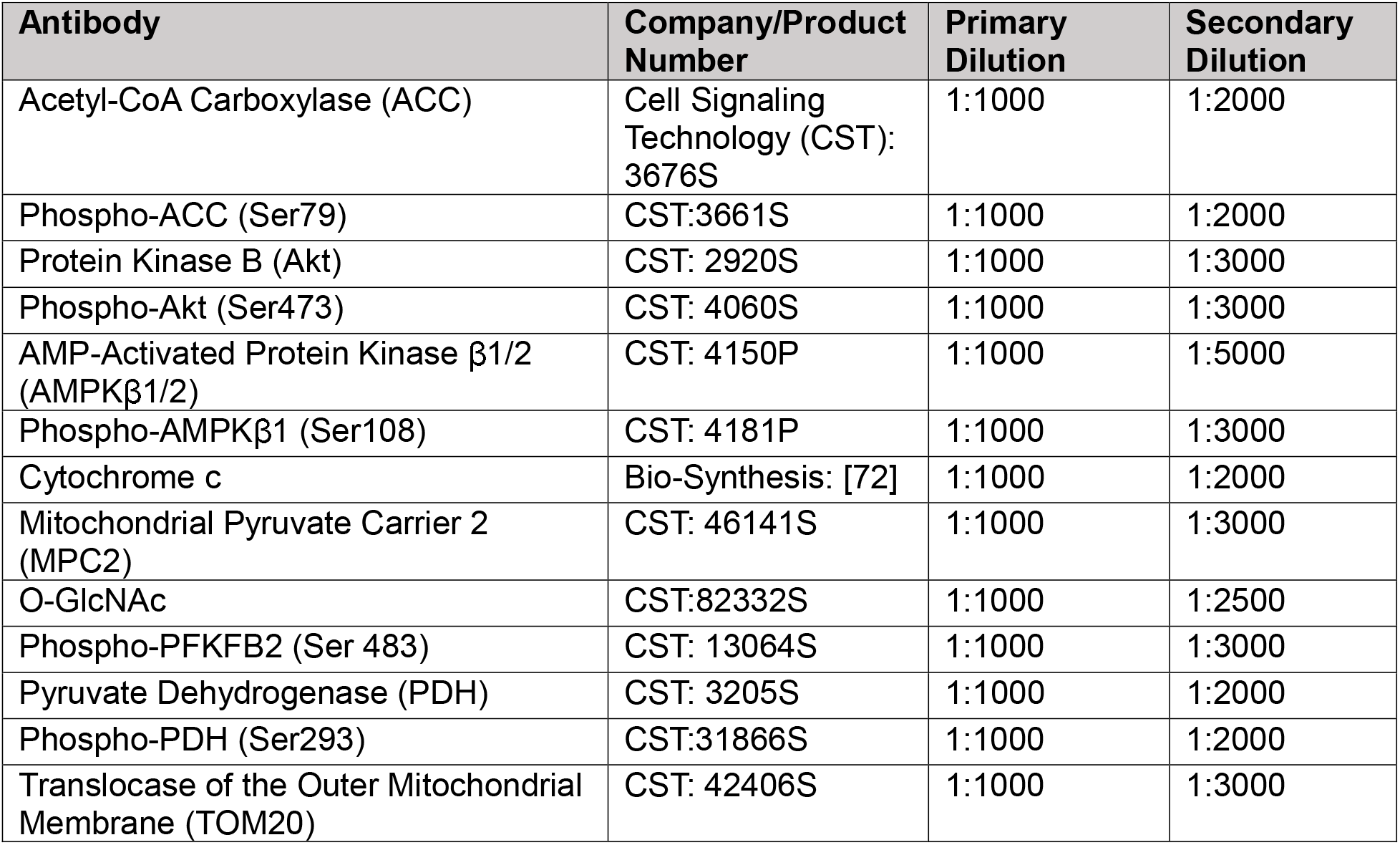
Antibodies used for Western Blot Analysis.

| Antibody | Company/Product Number | Primary Dilution | Secondary Dilution |
| --- | --- | --- | --- |
| Acetyl-CoA Carboxylase (ACC) | Cell Signaling Technology (CST): 3676S | 1:1000 | 1:2000 |
| Phospho-ACC (Ser79) | CST:3661S | 1:1000 | 1:2000 |
| Protein Kinase B (Akt) | CST: 2920S | 1:1000 | 1:3000 |
| Phospho-Akt (Ser473) | CST: 4060S | 1:1000 | 1:3000 |
| AMP-Activated Protein Kinase β1/2 (AMPKβ1/2) | CST: 4150P | 1:1000 | 1:5000 |
| Phospho-AMPKβ1 (Ser108) | CST: 4181P | 1:1000 | 1:3000 |
| Cytochrome c | Bio-Synthesis: [72] | 1:1000 | 1:2000 |
| Mitochondrial Pyruvate Carrier 2 (MPC2) | CST: 46141S | 1:1000 | 1:3000 |
| O-GlcNAc | CST:82332S | 1:1000 | 1:2500 |
| Phospho-PFKFB2 (Ser 483) | CST: 13064S | 1:1000 | 1:3000 |
| Pyruvate Dehydrogenase (PDH) | CST: 3205S | 1:1000 | 1:2000 |
| Phospho-PDH (Ser293) | CST:31866S | 1:1000 | 1:2000 |
| Translocase of the Outer Mitochondrial Membrane (TOM20) | CST: 42406S | 1:1000 | 1:3000 |

### RT-qPCR

RNA was extracted from 10-15 mg of powdered frozen heart tissue (RNeasy Fibrous Tissue Mini Kit, Qiagen) and used for cDNA synthesis (QuantiTect Reverse Transcription Kit, Qiagen). RT-qPCR was performed (QuantiTect SYBR Green PCR Kit, Qiagen) in duplicate. The standard ΔCT method was used for data analysis with ΔCT equal to the geomean of CTs of reference genes Rpl4, Eef1e1, and Tbp subtracted from the CT of the gene of interest. Primers were obtained from Integrated DNA Technologies and sequences are included in **Table 2**.

**Table 2.**
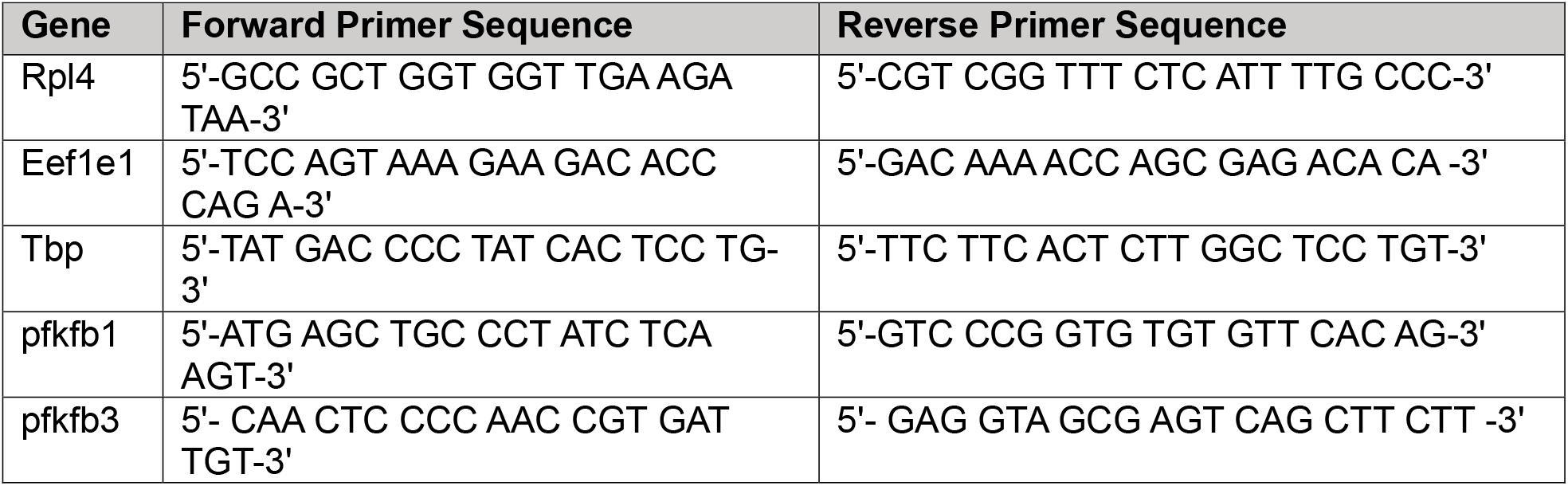
Primer sequences used for RT-qPCR.

### Electrocardiography

Surface electrocardiography (lead II) was recorded using three needle electrodes placed subcutaneously in the right front, right hind, and left hind limbs of the animal. General anesthesia was initially induced using 2.5% isoflurane and mice were maintained under 1.5% isoflurane during the recording process. Electrocardiographic signal was acquired using the PowerLab system (35 Series, AD Instruments) with amplification (FE231 BioAmp, AD Instruments). LabChart Software (Version 8, AD Instruments) was used for recording and analysis. A sampling rate of 4 kHz was utilized, with high and low pass filters set at 0.3 Hz and 1 kHz respectively. Data analysis was performed with blinding to genotypes. Beat averaging view was employed to measure wave amplitudes, segments, and interval durations in four beat average increments. The cumulative average of these four beat measurements over five minutes of recording was taken. However, due to artifactual isoelectric noise, elucidation of the exact peak and end of T waves was difficult in portions of the recordings. Thus, T waves were calculated from the average of 10 beats in the regions with lowest isoelectric noise. The corrected QT interval was calculated using the Bazett formula, 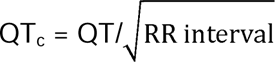

### Echocardiography

Echocardiographic measurement was performed on a cohort of male mice anesthetized using 2-2.5% isoflurane for induction and 1.0% isoflurane for maintenance. The Siemens Acuson CV-70 system with a 13 MHz probe was utilized for image acquisition and analysis. M-mode in parasternal long axis view and pulsed wave Doppler imaging in apical 4-chamber view were used to measure systolic and diastolic function respectively. Left ventricular mass was calculated using the Penn algorithm, 1.05*((diastolic interventricular septal wall thickness + diastolic left ventricular internal diameter + diastolic left ventricular posterior wall thickness)^3^ – (diastolic left ventricular internal diameter)^3^), as previously described [24, 25]. Myocardial performance index was calculated as (isovolumic contraction time + isovolumic relaxation time)/ejection time.

### Proteomics

Following homogenization and centrifugation at 550 x g, 100 µL of the supernatant was mixed with 100 µL 1% SDS and 20 µL 10% SDS, heated, and protein concentrations measured. Based on these concentrations, the volume containing 100 µg total protein was used for analysis and 100 µL 1% SDS was added, along with 1 µg BSA as a non-endogenous internal standard. The samples were mixed, heated at 70°C, and precipitated with acetone overnight. The precipitate was reconstituted in Laemmli sample buffer at 1 µg/µL and run 1.5 cm into an SDS-Page gel. Each 1.5 cm lane was cut as a complete sample, chopped into smaller pieces, washed, reduced, alkylated, and digested with 1 µg trypsin overnight at room temperature. Peptides were extracted from the gel in 50% acetonitrile. These extracts were dried by Speedvac and reconstituted in 200 µL 1% acetic acid for analysis.

For selected reaction monitoring (SRM), a ThermoScientific TSQ Quantiva instrument was used in SRM mode. Previously validated assay panels were used to measure the respective groups of proteins, with most assays monitoring two peptides per protein. A cycle time of 2 seconds was used to give approximately 15 data points across 30 second chromatographic peaks. The LC conditions were a linear gradient elution from 2%B to 50%B in 60 min.

For data independent acquisition (DIA), a ThermoScientific QEx plus instrument was used in the DIA mode with a 20 m/z window working from m/z 350 to 950. The orbitrap was operated at a resolution of 17,500. A full scan spectrum at a resolution of 70,000 was acquired each cycle. These conditions give 7-8 data points across our typical 30 second chromatographic peaks. DIA data were analyzed using the program Skyline based on a large group of internally developed assays, with each using 2-3 peptides, that have been validated in prior experiments.

For both SRM and DIA analyses, Skyline is used to locate and integrate the proper chromatographic peaks. Proper retention times are predicted based on retention time calibration using BSA and trypsin peptides with manual inspection and adjustment as needed. Calculations determine the total protein response from the geomean of the two monitored peptides. Results are normalized to the BSA internal standard and expressed as pmol/100 µg total protein.

### Metabolomic analysis using LC-MS & GC-MS

LC-MS targeted and GC-MS semi-targeted metabolomic analyses were performed from a single preparation. Briefly, frozen heart tissue was pulverized, followed by metabolite extraction with methanol:water (8:2 v/v), sonication, and centrifugation. Five µl of adonitol (1 mg/ml) and 5 µl of ^13^C_3_ lactate (0.5 mg/ml) were added as internal standards to samples. Metabolite extracts were divided into 2 aliquots, with one used for GC-MS analysis, and the other used for LC-MS analysis. Pooled samples, prepared by combining supernatants from every sample, were used as quality control (QC) samples during the analysis.

For LC-MS targeted analysis, supernatant was dried and reconstituted in 100µl of 7:3 v/v acetonitrile and water and transferred to glass inserts for LC-MS analysis on Agilent 6546 LC/Q- TOF,- coupled to an Agilent 1290 Infinity II LC. Chromatographic separation was performed on an Agilent InfinityLab Poroshell 120 HILIC-Z, 2.1 × 150 mm, 2.7 μm column, coupled with a UHPLC Guard, HILIC-Z, 2.1 mm × 5 mm, 2.7 μm, at 15°C, with a total run time of 29 min. The mobile phase consisted of 20 mM ammonium acetate buffer in water (pH=9.2) (A) and acetonitrile (B). To ensure a constant concentration during gradient elution, the InfinityLab deactivator additive (p/n 5191-4506) was added to the aqueous mobile phase. A linear gradient elution was used at a flow rate of 0.4 ml/min with the following program: 0.0 min (15% A and 85% B), 1.0 min (15% A and 85% B), 8 min (25% A and 75% B), 12.0 min (40% A and 60% B), 15.0 min (80% A and 20% B), 18.0 min (80% A and 20% B), 19.0 min (15% A and 85% B) and 29.0 min (15% A and 85% B). Data was acquired in a negative ESI full MS scan mode (scan range: m/z 40 to 1,000) using Agilent MassHunter Acquisition software version 10.0. Optimum values for MS parameters were as follows: gas temperature 300°C; drying gas flow 13 L/min; Nebulizer pressure 40 psi; Sheath gas temperature 350°C; Sheath gas flow 12 L/min; Capillary voltage 3500 V; Nozzle voltage 0 V; skimmer offset 45 V; Fragmentor 125 V; Octopole 1 RF Voltage 750 V.

Data analysis and peak integration for target metabolites was performed with Agilent MassHunter Quantitative analysis software (version 10.1). An analytical standard mix of glycolytic and TCA cycle intermediates was analyzed along with samples for identification of these metabolites in the samples. A custom Agilent Personal Compound Database and Library (PCDL) of target metabolites (glycolytic and tricarboxylic acid cycle intermediates, amino acids and nicotinamide adenine dinucleotide reduced (NADH) and oxidized (NAD^+^) forms) was created from Agilent METLIN PCDL for identification within the samples. The retention times derived from analyses of an in-house prepared mix of analytical glycolytic and tricarboxylic acid cycle intermediate standards, a canonical amino acid unlabeled standard mix (Cambridge isotope laboratories, Inc., Cat. No. MSK-CAA-US-1), and a NAD+ standard were also included in the custom PCDL to facilitate the identification of metabolites in the samples. Relative abundance values for target metabolites were obtained by normalizing the raw data with tissue weights and adonitol (added as internal standard) values.

For GC-MS analysis, supernatant was dried & derivatized in two steps and the analysis was performed as previously published using GC-MS (Agilent 7890B-5977A) [26]. Briefly, samples were dissolved in 25 μL of 20 mg/mL methoxyamine hydrochloride in pyridine for 1.5 hours at 37°C with constant orbital shaking. Second, 35 μL Bis(trimethylsilyl)trifluoroacetamide (BSTFA) was added, and the samples were mixed at 37°C for 40 minutes.

After derivatization, samples were transferred to glass capped vials and analyzed on the GC-MS system with EI source in a scan mode (70-600 m/z). An alkane mix (C10 to C24), an analytical standard mix of glycolytic and TCA cycle intermediates, and a canonical amino acid unlabeled standard mix (Cambridge isotope laboratories, Inc., Cat. No. MSK-CAA-US-1) were prepared in the same manner as samples and analyzed along with samples. Alkane mix was used as a retention time standard ladder. The analytical standard mix prepared from standards of pyruvate, lactate, succinic acid, fumaric acid, malic acid, aspartic acid, glutamic acid, citric acid, glucose, fructose-6-phosphate, and glucose-6-phosphate was used for confirmation of these metabolites in the samples. Similarly, analysis of an amino acid standard mix was used for confirmation of amino acids in the samples. Data was processed using Mass Hunter Quantitative data analysis software with the integrated mass-spectrometry library from the National Institute of Standards and Technology (NIST, Gaithersburg). The raw data were normalized by tissue weight and adonitol response. All raw GC–MS data is available upon request. All chemicals and analytical standards (except for the amino acid standard mix) were purchased from Sigma Aldrich (St. Louis, MO, USA) with exception of the pyridine and methoxyamine hydrochloride (Thermo Fisher Scientific, Waltham, MA, USA).

### Pathway Analysis and Principal Component Analysis

Integration of LC and GC mass spectrometry data was performed by Z-standardization. The joint data were then used for principal component analysis (PCA). To generate a heat map, ratios of metabolite abundances in cKO relative to CON animals were first logarithmically transformed. Significantly different metabolites in cKO relative to CON animals were identified using Student’s t-test with p-value threshold at 0.05 and subjected to pathway analysis using the MetaboAnalyst online tool [27] on a KEGG background [28]. Similarly, proteomics data were Log_10_-transformed prior to Student’s t-test to identify proteins with significantly altered abundance. Ratios of protein abundance in cKO animals relative to CON were used to build a heatmap. Upregulated and downregulated enzymes were subjected to proteomic pathway analysis separately using the STRING online tool [29] on a KEGG background [28]. The pathways are presented according to false discovery rate (FDR)-corrected q-values <0.05 of association significance.

### Statistics

Data were analyzed using a two-tailed, unpaired Student’s *t* test using GraphPad Prism (Version 10) and presented as mean ± SD unless otherwise indicated. Statistical significance was established at p≤0.05.

## Results

### Validation of PFKFB2 knockout mice and absence of compensation by PFK-2 isoforms

*Pfkfb2^fl/fl^* mice were crossed with heterozygous cardiomyocyte-specific Cre expressing *Pfkfb2^fl/fl^* mice to generate a mix of *Pfkfb2^fl/fl^ Cre-* (CON) and *Pfkfb2^fl/fl^ Cre+* (cKO) mice. Mass spectrometry confirmed that PFKFB2 was undetectable in cKO animals (Figure 1A). Further, western blot analysis using an antibody against phospho-PFKFB2 (Ser 483) also showed no detectable signal in cKO hearts (Figure 1B). To determine whether the loss of PFKFB2 promoted compensatory upregulation of other PFK-2 isoforms, RT-qPCR was performed. Gene expression data shows no transcriptional upregulation of *Pfkfb1* and *Pfkfb3* isoforms in cKO hearts (Figure 1C). Interestingly, *Pfkfb1* transcript expression was reduced by 82.5% (p=0.0032, Figure 1C).

**Figure 1.**
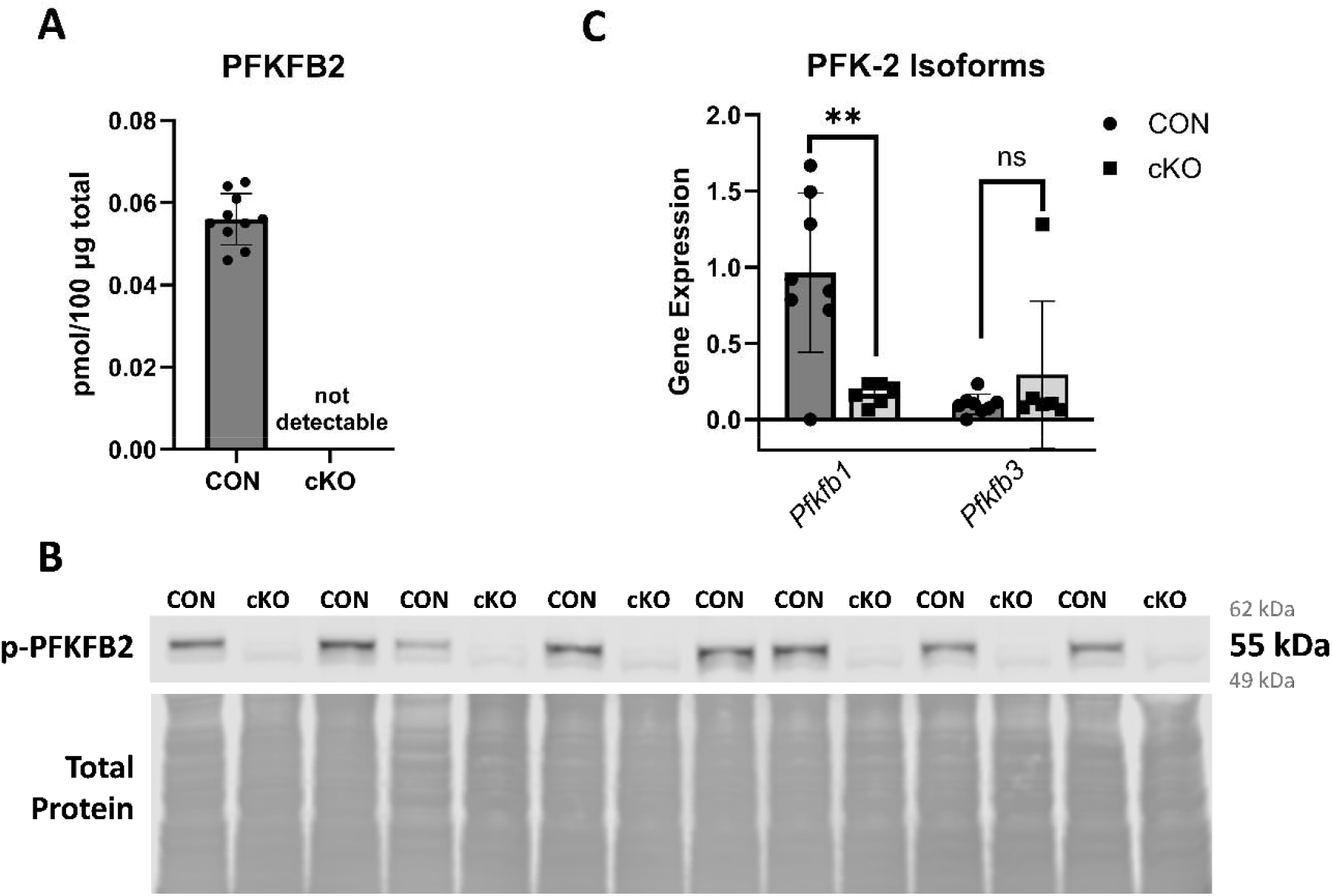
Validation of *Pfkfb2* knockout and absence of compensation by other isoforms. **A**, PFKFB2 is detectable by mass spectrometry in control (CON; n=10) but not *Pfkfb2* knockout (cKO; n=8) animals. **B**, p-PFKFB2 (Ser 483) is detectable in CON (n=8) but not cKO (n=6) animals by western blot. **C**, No compensation by *Pfkfb1 or Pfkfb3* isoforms is observed in cKO (n=6) animals relative to CON (n=8) as measured by RT-qPCR. Data are shown as mean ± SD. ** p ≤ 0.01, unpaired Student’s *t* test.

We also monitored basic physiological parameters to determine potential systemic effects of the *Pfkfb2* knockout. No differences were observed in body weight normalized to tibia length or blood glucose levels in cKO animals relative to CON (**Table 3**). However, we did observe a modest but significant 20.4% increase in heart weight (p=0.0006; **Table 3**), implying potential pathology. Notably, these animals also exhibited shortened lifespans, with sudden unexplained mortality at approximately 9 months of age without prior signs of progressive decline such as alterations in body weight or distressed appearance. Further metabolic and physiological characterization sought to elucidate underlying causes of this sudden mortality.

**Table 3.**
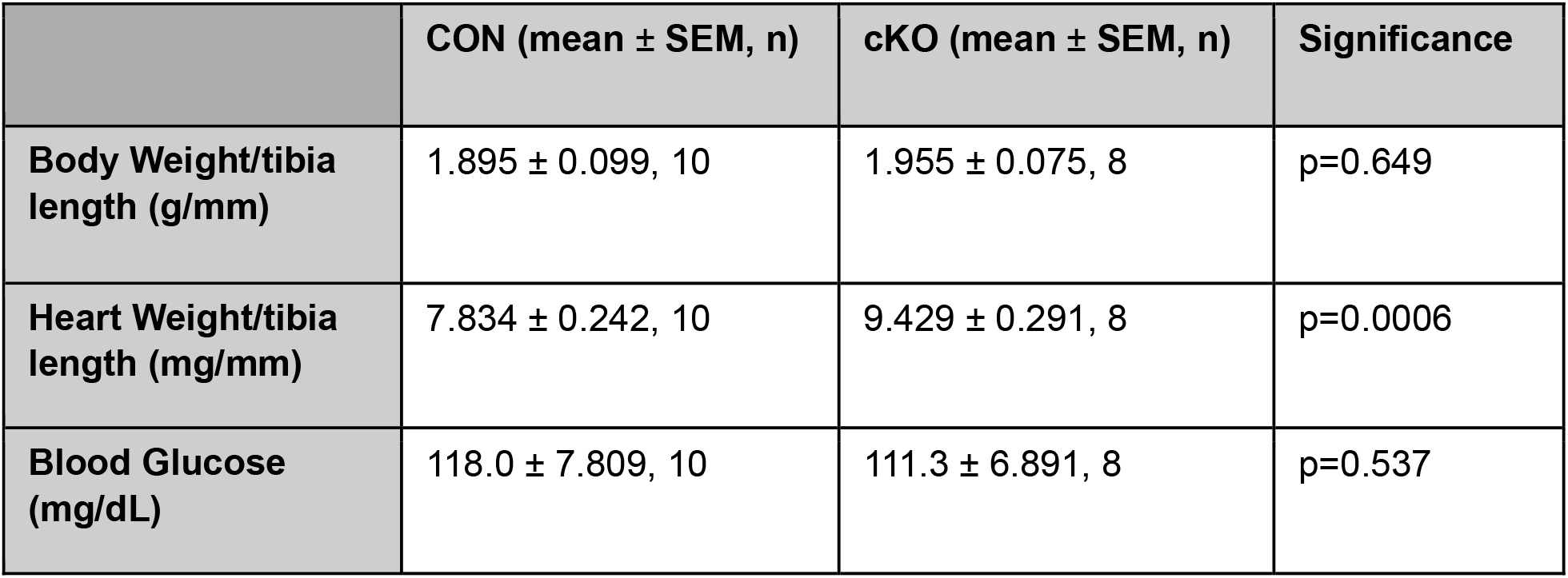
Systemic impacts of *Pfkfb2* are minimal. *Data were analyzed by unpaired Student’s *t* test.

| | CON (mean ± SEM, n) | cKO (mean ± SEM, n) | Significance |
| --- | --- | --- | --- |
| Body Weight/tibia length (g/mm) | 1.895 ± 0.099, 10 | 1.955 ± 0.075, 8 | p=0.649 |
| Heart Weight/tibia length (mg/mm) | 7.834 ± 0.242, 10 | 9.429 ± 0.291, 8 | p=0.0006 |
| Blood Glucose (mg/dL) | 118.0 ± 7.809, 10 | 111.3 ± 6.891, 8 | p=0.537 |

### Metabolomic and proteomic analyses suggest metabolic reprogramming

To determine the metabolic effects of *Pfkfb2* knockout, semi-targeted metabolomic analyses using both LC and GC mass spectrometry were performed. Fifty-four metabolites were identified using this approach (select metabolites shown in Figures S1 and S2). Principal component analysis revealed a separation of CON and cKO groups (Figure 2A), suggesting that knocking out *Pfkfb2* induces broad changes in cardiac metabolism. A heatmap of differences in metabolites in cKO relative to CON hearts was generated (Figure 2B), with metabolites that were significantly different indicated in bold (18 of the 54 total). Pathway analysis of these significantly different metabolites (Figure S2T) indicated the largest effect was on amino acid metabolism pathways (Figure S2A-S), with an additional notable impact on nicotine and nicotinamide metabolism driven by decreased abundance of NAD^+^ (Figure S1T) and its precursor nicotinamide (Figure S1U). Somewhat unexpectedly, knocking out *Pfkfb2* had minimal effect on glucose or glycolytic intermediates (glucose-6-phosphate, fructose-6-phosphate, fructose-1,6-bisphosphate, dihydroxyacetone phosphate, glyceraldehyde-3-phosphate, 3-phosphoglyceric acid, and phosphoenolpyruvate), except for a significant increase in pyruvate (Figure S1A-I). However, several glucose-derived metabolites, including ascorbic acid, aspartic acid, serine, glycine, and glutamic acid were significantly increased in cKO hearts (Figures S1V, S2D-E, S2G, S2O).

**Figure 2.**
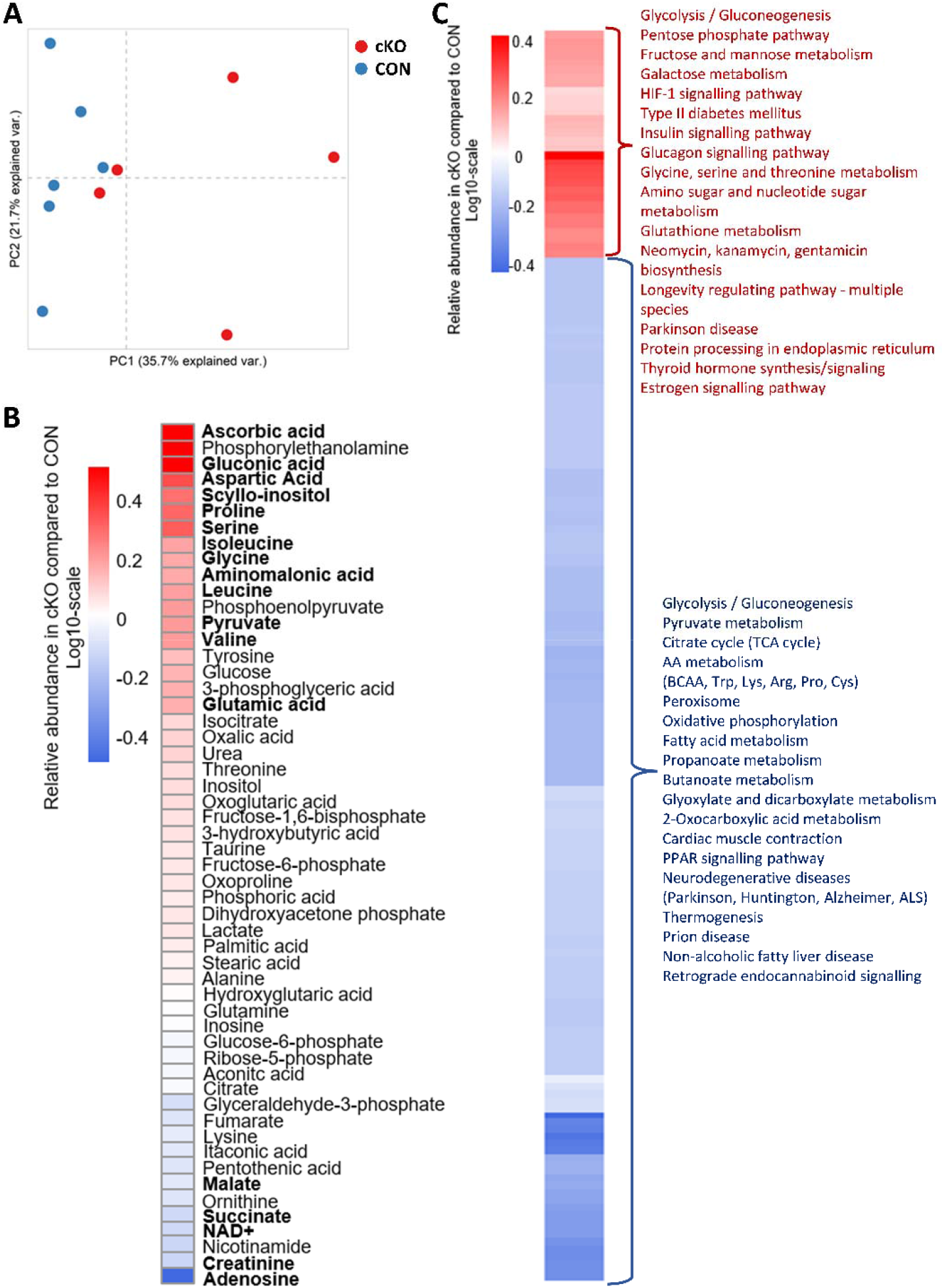
*Pfkfb2* knockout drives metabolic reprogramming. **A**, Principal component analysis of metabolites measured using a combination of liquid chromatography-mass spectrometry (LC-MS) and gas chromatography-mass spectrometry (GC-MS) demonstrating separation of *Pfkfb2* knockout (cKO) and control (CON) animals. **B**, Heat map of metabolites from LC-MS and GC-MS generated via logarithmic transformation of the ratio of metabolite abundance in cKO relative to CON hearts. An unpaired Student’s *t* test was used and resulting significant metabolites are bolded. **C**, Ratios of protein abundances in cKO relative to CON animals were logarithmically transformed and used to generate a heat map. A Student’s *t* test was then applied and significantly different proteins were subjected to pathway analysis. Upregulated pathways are listed in red and downregulated pathways are listed in blue.

Proteomics was next conducted to determine how the absence of PFKFB2 affects metabolic reprogramming. Despite the lack of differences in glycolytic intermediates, at the protein level there were significant increases in the insulin-independent glucose transporter GLUT1 (Figure S3A) and glycolytic enzymes (Figure S3C-L). The increase in glycolytic enzymes occurred with a universal decrease in abundance of mitochondrial proteins of the TCA cycle (Figure S4) and electron transport chain (Figures S5 and S6) in cKO animals relative to CON. These changes are indicative of a decrease in mitochondrial abundance, and it is therefore unsurprising that proteins involved with mitochondria-localized metabolic pathways, such as beta-oxidation, are also decreased (Figure S7A-G). Though, decreased mitochondrial beta oxidation occurred in conjunction with commensurate reductions in enzymes of peroxisomal beta oxidation (Figure S7G-K) and other aspects of fatty acid metabolism (Figure S8) in cKO animals. Also striking were reductions in proteins involved with amino acid catabolism (Figure S9), which could underlie the general increase in amino acid abundance observed by metabolomic analyses.

Stress response pathways were also different in cKO animals. Proteins responsible for maintenance of redox balance were upregulated (Figure S10A-B, F-G). Similarly, increases in chaperone proteins including DJ-1 (parkinsonism associated deglycase), heat shock proteins, and protein disulphide isomerase A3 were also observed (Figure S10H-K). However, predominantly mitochondria-localized proteins in these classes, e.g. superoxide dismutase or prohibitin, were decreased or unchanged (Figure S10D, L), consistent with decreased mitochondrial abundance in cKO animals.

Proteomics data also suggests potential for proteostatic alterations in cKO animals. There is an increase in eukaryotic translation elongation factor that points towards increased protein synthesis (Figure S11A). Reciprocally, the upregulation in ubiquitin B, ubiquitin-like modifier activating enzyme 1, valosin containing protein (p97), lon peptidases, and vacuolar protein sorting 35 suggest the potential for increased protein degradation (Figure S11B-F).

Pathway analysis of proteomics data highlighted many of these changes (Figure 2C). Glycolysis and HIF-1 signaling were among the most prevalent upregulated pathways in the cKO heart. Oxidative phosphorylation and the TCA cycle were identified as being downregulated, which is consistent with a decreased mitochondrial abundance in cKO hearts. These changes culminate in a phenotype indicative of a metabolic shift to glucose oxidation in the absence of PFKFB2.

### Metabolic Kinase-driven shift toward glucose oxidation

We further characterized the metabolic phenotype of cKO hearts by examining the protein levels of AMPK and Akt. These two kinases serve as metabolic regulators in response to extracellular cues, nutrient availability, and stresses of metabolic and non-metabolic nature. We observed a significant increase in the expression of AMPKβ1/2 isoforms (49.7%; p=0.0006; Figure 3A). While the increase in the AMPKβ1 isoform alone failed to reach significance, we observed an increase in its activated form, p-AMPKβ1 (Ser108) (64.5%; p=0.0013; Figure 3A) in cKO animals. However, the ratio of p-AMPKβ1/AMPKβ1 was not different between cKO and CON hearts (Figure 3A). We next looked at the total levels of acetyl CoA carboxylase (ACC) and its phosphorylation status at Ser 79, a canonical substrate site for AMPK. Phosphorylation of ACC inhibits its activity, decreasing malonyl-CoA production, and promotes fatty acid oxidation (FAO). Interestingly, as with AMPKβ1, content of both phosphorylated and total ACC was increased (Figure S12) such that ratio between p-ACC and total ACC was not different between cKO and CON. While this supports that there is increased AMPK signaling, conclusions regarding malonyl-CoA production and regulation of FAO via ACC activity are ambiguous.

**Figure 3.**
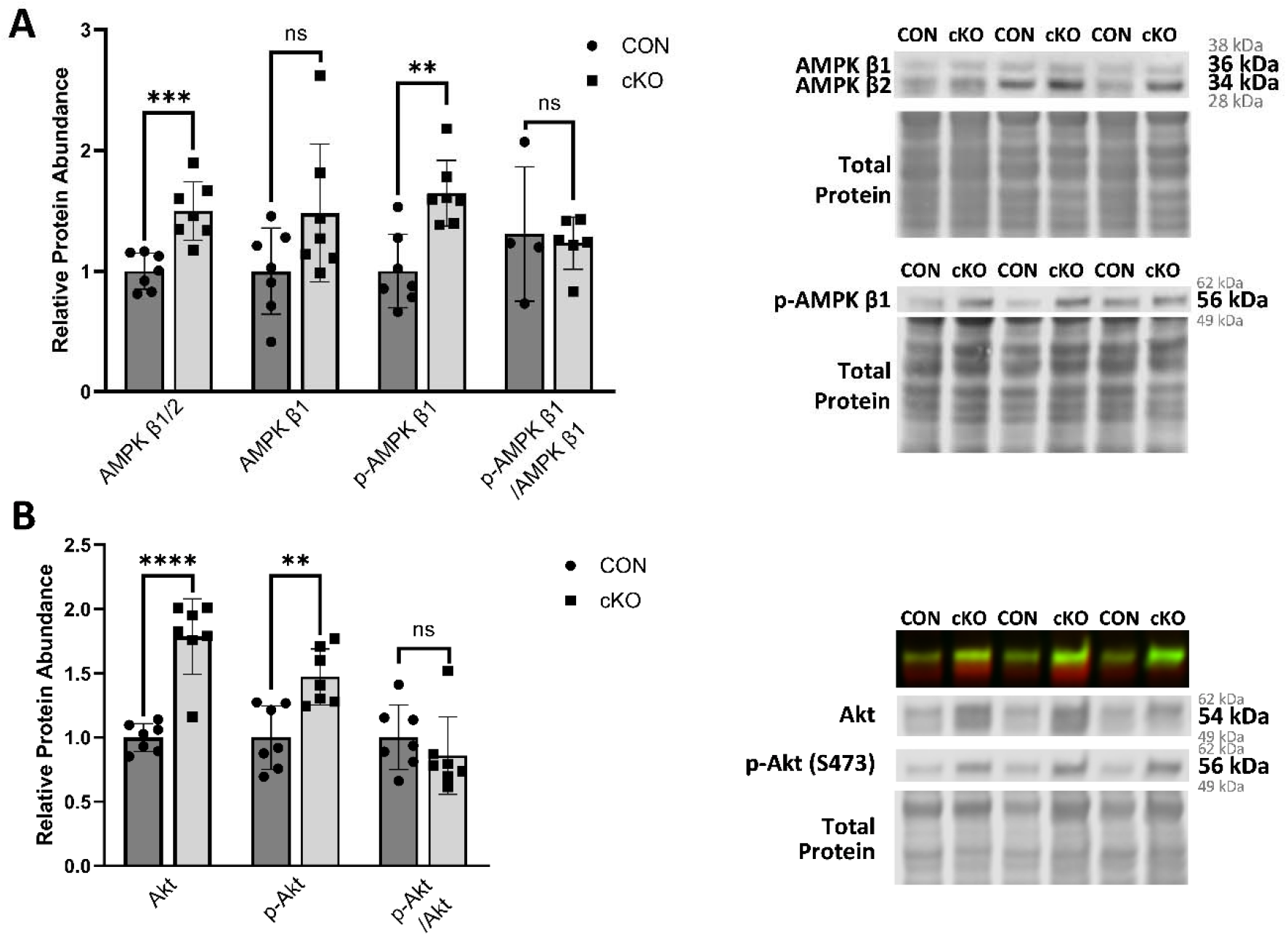
Abundance of metabolic kinases is increased with *Pfkfb2* knockout. **A**, AMPK β1/2, AMPK β1, and p-AMPK β1 (Ser108) were measured by western blot in knockout (cKO; n=7) relative to control (CON; n=7) animals and the ratio of p-AMPK β1/AMPK β1 was calculated (n=4-6). **B**, Akt and p-Akt (Ser473) were measured by western blot and the ratio of p-Akt/Akt was calculated in cKO (n=7) relative to CON (n=7). Data are shown as mean ± SD. Not significant (ns): p>0.05, **p≤0.01, ***p≤0.001, ****p≤0.0001, unpaired Student’s *t* test.

As with AMPKβ1, abundances of Akt and p-Akt were increased 78.6% (p<0.0001) and 47.2% (p=0.0025) respectively without a difference in the p-Akt to Akt ratio in cKO animals relative to CON (Figure 3B). The phosphorylated forms of these stress-activated kinases are generally associated with increased glucose uptake and oxidation through PFKFB2-dependent and independent means [2, 30-32]. Among these mechanisms, chronic activation of Akt can promote transcription of the gene encoding GLUT-1 [33], which may contribute to the upregulation found in cKO animals (Figure S3A).

We next performed functional assays to determine if knocking out PFKFB2 resulted in changes to PDH activity and substrate-specific mitochondrial respiration. These assays can distinguish between mitochondria that are primarily fatty acid-oxidation or glucose oxidation driven [18, 34, 35]. Respirometry with pyruvate (Pyr) and malate as substrates exhibited a trend toward greater ADP-dependent (state 3) respiration in cKO relative to CON hearts, though this difference failed to reach statistical significance (p=0.068; Figure 4A). Furthermore, no difference was observed in palmitoyl carnitine (PC) and malate-supported state 3 respiration (Figure 4B). However, the ratio of PC to Pyr state 3 respiration rates was significantly decreased in cKO mice (13.5%; p=0.002; Figure 4C), demonstrating that cKO heart mitochondria have greater maximal respiration rates with Pyr relative to PC as compared to CON. PDH activity measurements support these data. Basal PDH activity was increased 2.80-fold in cKO mitochondria relative to CON (p=0.001; Figure 4D). Interestingly, PDH from cKO animals did not respond to activation with Pyr and malate (Figure 4E), suggesting near maximal activity at baseline. This increased activity is mediated in part by a significant increase in PDH expression (24.1%; p=0.013) and a 15.7% decrease in inhibitory PDH phosphorylation at Ser293 (p=0.039; Figure 4F) [36]. The ratio of p-PDH to PDH was 31.0% less in cKO hearts as compared to CON (p=0.0063; Figure 4F). This is likely attributable in part to decreased abundance of pyruvate dehydrogenase kinase (PDK) isoforms (23.5-42.1%; p=0.0004-0.0026; Figure 4G), responsible for inhibitory PDH phosphorylation [37, 38].

**Figure 4.**
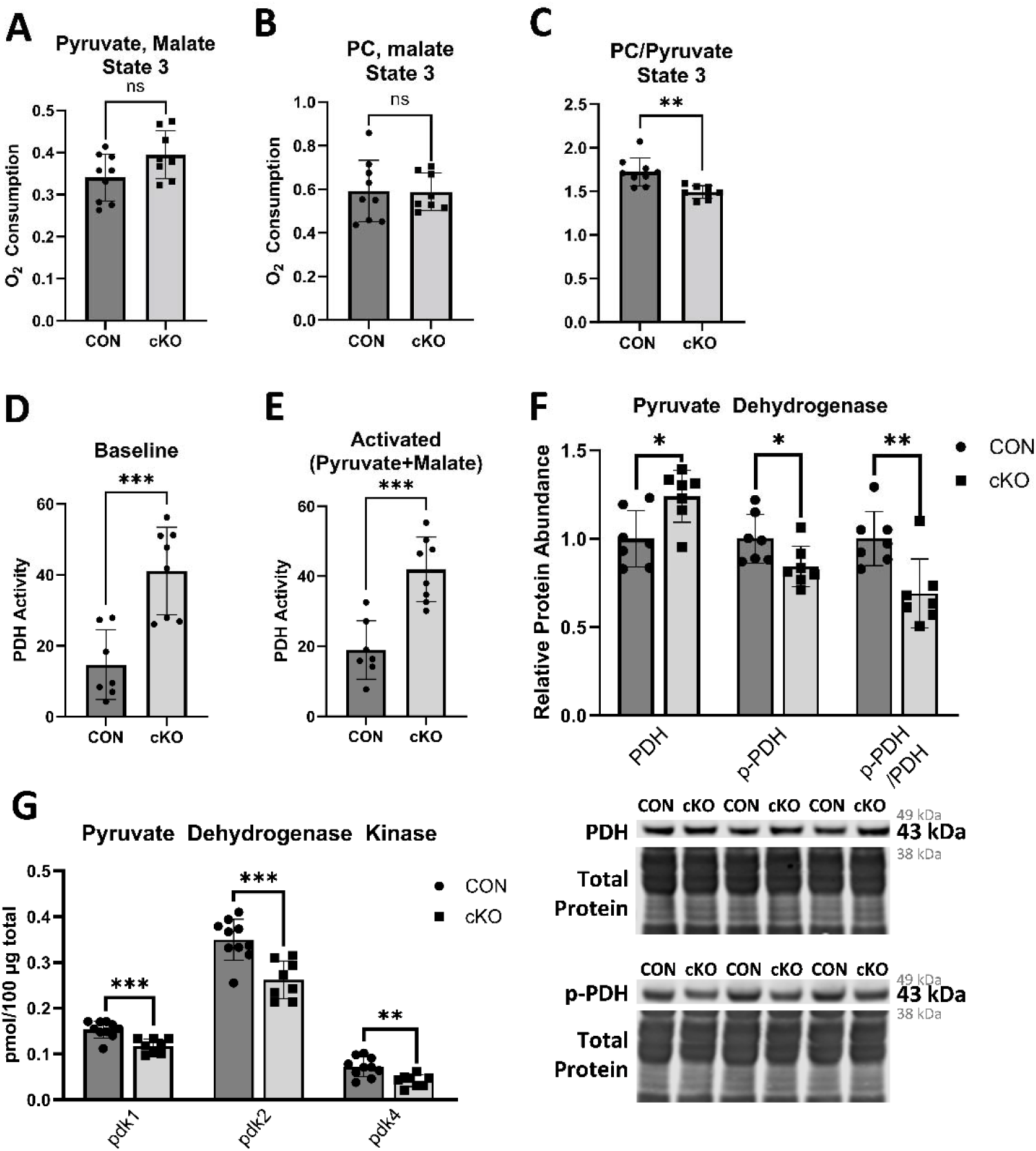
Loss of PFKFB2 promotes a shift toward pyruvate dependent respiration. **A-B**, Mitochondrial state 3 respiration as stimulated by pyruvate and malate or palmitoyl carnitine (PC) and malate respectively in *Pfkfb2* knockout (cKO; n=8) relative to control (CON; n=9) animals. **C**, The ratio of PC and malate-stimulated to pyruvate and malate-stimulated state 3 respiration in cKO (n=8) compared to CON (n=9) animals. **D**, Baseline pyruvate dehydrogenase (PDH) activity in cKO (n=8) relative to CON (n=7). **E**, Activated pyruvate dehydrogenase activity following stimulation with pyruvate and malate in cKO (n=8) relative to CON (n=7). **F**, Relative abundances of PDH, p-PDH (Ser 293), and the ratio of phosphorylated to total PDH in cKO (n=7) relative to CON (n=7) animals as measured by western blot. **G**, Relative abundance of pyruvate dehydrogenase kinase isoforms in cKO (n=8) relative to CON (n=10) animals measured by mass spectrometry. Data are shown as mean ± SD. Not significant (ns): p>0.05, *p ≤ 0.05, **p≤0.01, ***p≤0.001, unpaired Student’s *t* test.

The regulation of PDH activity is complex. PDH is primarily regulated by pyruvate dehydrogenase kinase and phosphatase activities, which in turn are regulated by acetyl-CoA levels and rapid turnover [37]. A previous study has also shown that mitochondria-associated Akt can relay cellular signals to activate PDH [39]. Thus, we investigated mitochondrial Akt abundance and sublocalization within the mitochondria. Possibly contributing to the observed increase in PDH activity, both total Akt and phosphorylated Akt (Ser473) were increased by over 3-fold in the mitochondrial fractions of cKO relative to CON animals (p=0.0006-0.0026; Figure S13A). To determine sublocalization of Akt and p-Akt, isolated mitochondria were treated with proteinase K in the presence or absence of Triton X-100. Proteins degraded in the absence of Triton X-100 are localized to the outer mitochondrial membrane; proteins degraded only in the presence of Triton X-100 are either in the intermembrane space, inner membrane, or matrix. PDH, MPC2, cytochrome c, and TOM20 were used as matrix, mitochondrial inner membrane, intermembrane space, and outer membrane controls. As expected, PDH, MPC2, and cytochrome c were protected from proteinase K degradation in the absence of Triton X-100 (Figure S13B). However, TOM20 and the majority of Akt (84%) and p-Akt (97%) were degraded despite Triton X-100 absence (Figure S13B). This suggests that while a portion of mitochondria-associated Akt may be found inside the mitochondria, a majority is likely localized to the outer mitochondrial membrane. Cumulatively, respirometry data, PDH activity, proteomics findings, and greater mitochondria associated Akt are suggestive of preferential glucose oxidation and diminished FAO in cKO animals.

### Metabolite shunting to ancillary glucose metabolism pathways

Previous studies from the Hill lab have shown that PFK-2 activity is a key regulator of the metabolic fate of glucose. Activation of PFK-2 couples increased glucose oxidation to concomitant glycolytic upregulation, thereby mitigating flux through ancillary glucose metabolic pathways [40]. We therefore examined ancillary pathways to see if there are indications of increased glucose metabolism via auxiliary routes in cKO hearts. The pentose phosphate pathway (PPP) was determined by proteomic-based pathway analysis to be upregulated in cKO animals relative to CON (Figure 2C). This is partially attributable to upregulations in key enzymes of the oxidative branch of the pathway, glucose-6-phosphate dehydrogenase (76.3%; p<0.0001; Figure 5) and 6-phosphogluconate dehydrogenase (74.5%; p<0.0001; Figure 5). However, no accumulation of the PPP product ribose-5-phosphate is observed in cKO animals (Figure 5). This is likely due to a concordant increase in the downstream non-oxidative branch of the PPP, as indicated by a modest increase in transaldolase (24.9%; p=0.0238; Figure 5) and a prominent increase in transketolase (91.0%; p<0.0001; Figure 5). These changes in enzyme abundance are consistent with shunting of glucose-6-phosphate to the oxidative PPP branch, followed by a single- or multi-step conversion of ribose-5-phosphate to glyceraldehyde-3-phosphate through non-oxidative processes [41]. This would ultimately support pyruvate production via glycolytic reactions downstream of glyceraldehyde-3-phosphate, bypassing the PFK-2/PFK-1 regulatory nexus. Of additional relevance, the oxidative branch of the PPP produces cytosolic NADPH [42]. Interestingly, NADPH-dependent antioxidant enzymes including thioredoxin, thioredoxin reductase, glutathione peroxidase, and glutathione disulfide reductase, are increased in cKO animals (23.3-86.8%; p≤0.0007; Figure S10A-B, F-G). Thus, PPP upregulation may have implications beyond auxiliary support of glucose oxidation.

**Figure 5.**
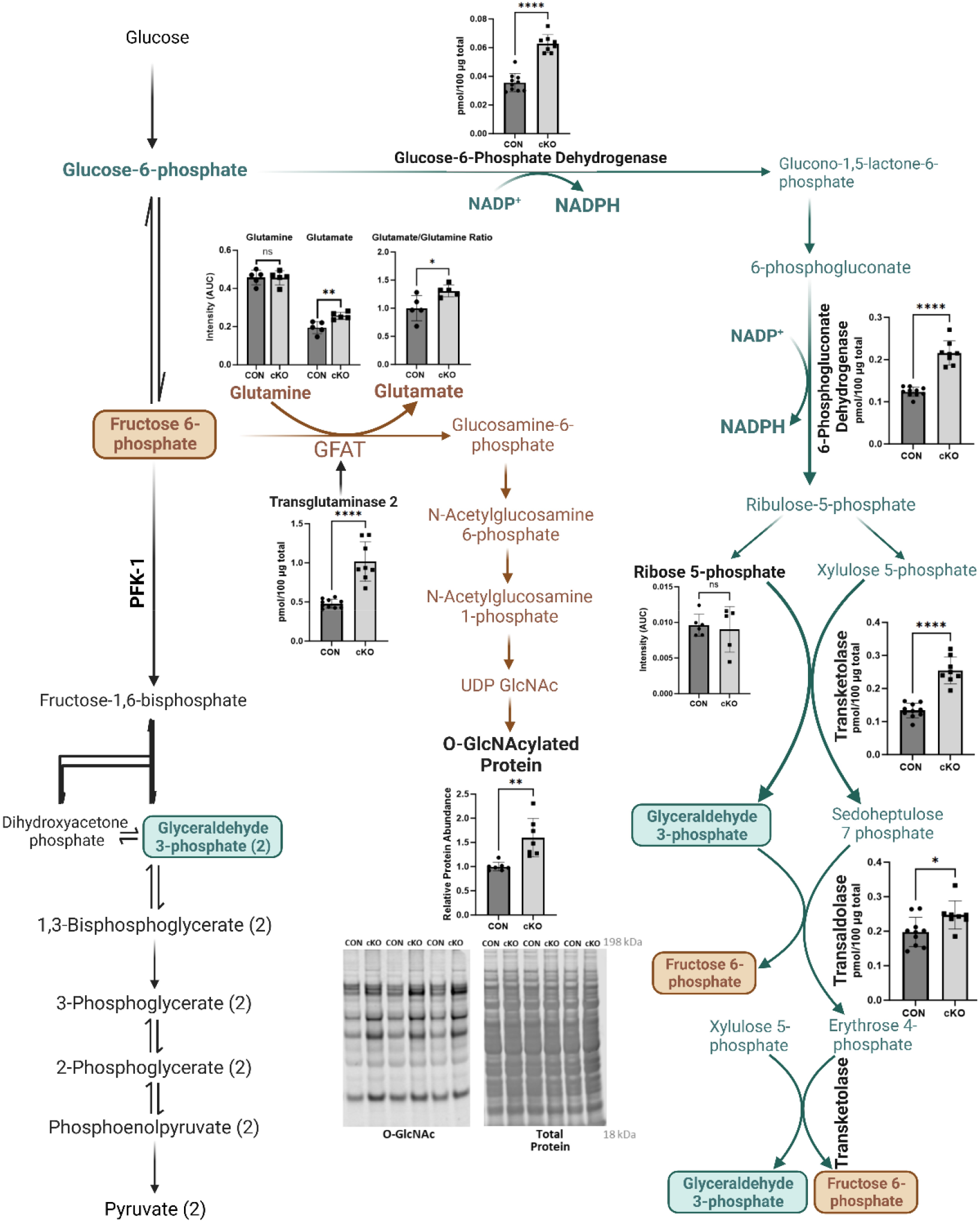
Ancillary pathways are activated in *Pfkfb2* knockout (cKO) animals relative to controls (CON). Glucose 6 phosphate dehydrogenase, 6-phosphonogluconolactone dehydrogenase, transaldolase, and transketolase were measured using mass spectrometry in cKO (n=8) and CON (n=10) animals. Glutamate and Glutamine were measured using liquid chromatography mass spectrometry in cKO (n=5) and CON (n=5) animals. Ribose 5 phosphate was measured using gas chromatography mass spectrometry in cKO (n=5) and CON (n=6) animals. The post-translational modification O-GlcNAcylation was measured as a proxy for hexosamine biosynthesis activity using western blot with quantification of the full lane (n=7/group). Data are shown as mean ± SD. Not significant (ns): p>0.05, * p ≤ 0.05, **p≤0.01, ****p≤0.0001, unpaired Student’s *t* test. Figure was made using BioRender.com.

The PPP is not the sole branch of glycolysis utilizing metabolites upstream of PFK-1 that is upregulated. The pathway responsible for ascorbic acid biosynthesis in mice, and most non-human mammals, shunts glucose-6-phosphate from glycolysis [43]. An upregulation of this process in cKO animals is suggested by a 6.58-fold greater ascorbic acid abundance (Figure S1V). The HBP is another ancillary pathway of importance as it utilizes fructose-6-phosphate, the metabolite directly upstream of PFK-1 in glycolysis. HBP activity can be measured by assessing O-GlcNAcylation levels of proteins, as this posttranslational modification depends on the HBP endpoint as a substrate [44]. Consistent with the notion that glucose is being rerouted upstream of the PFK-2/PFK-1 regulatory nexus, we observed a 59.9% increase in O-GlcNAcylation (Figure 5; p=0.0021) in cKO hearts. Increased HBP activity is further supported by an increase in glutamate abundance, thus increasing the glutamate/glutamine ratio (Figure 5), as glutamine is converted to glutamate in the HBP [45]. Additionally, abundance of transglutaminase 2, a protein that stimulates the hexosamine biosynthesis pathway through activation of its first and rate limiting enzyme, glucosamine:fructose-6-phosphate amidotransferase (GFAT) [46], is increased by over 2-fold (p<0.0001; Figure 5). O-GlcNAclylated proteins play critical roles in physiology and pathophysiology of the heart and thus, an increase in HBP activity has significant implications [16, 47].

### Structural and functional changes in PFKFB2 cKO hearts

Chronic HBP activation and other aspects of metabolic reprogramming can lead to eccentric left ventricular (LV) remodeling [48]. This can manifest as LV dilation and concomitant systolic dysfunction, which was indeed observed by echocardiography in cKO mice. LV dilation, as indicated by increased LV internal diameter at diastole (22.6%, p=0.004; **Table 4**), occurs without consequent left ventricular wall thinning in cKO hearts, culminating in an increase in left ventricular mass (30.7%, p=0.016; **Table 4**). Corroborating this, heart weight normalized to tibia length increased (**Table 3**) in cKO relative to CON animals, while diastolic left ventricular posterior wall thickness (**Table 4**) and interventricular septal wall thickness (**Table 4**) were unchanged. In addition to these structural changes, systolic function was impaired. This was indicated by 35.1% decreased fractional shortening (FS; p=0.0004; **Table 4**) in cKO animals. Though still preserved, ejection fraction (EF) was decreased by 23.7% in cKO animals relative to CON (p=0.0002; **Table 4**).

**Table 4.** Echocardiographic remodeling occurs in response to *Pfkfb2* knockout. *Data were analyzed by unpaired Student’s *t* test.

| | CON (mean ± SEM, n) | cKO (mean ± SEM, n) | Significance |
| --- | --- | --- | --- |
| Left Ventricular Internal Diameter, Diastolic (mm) | 3.671 ± 0.109, 7 | 4.500 ± 0.212, 4 | p = 0.004 |
| Left Ventricular Posterior Wall Thickness, Diastolic (mm) | 0.914 ± 0.026, 7 | 0.850 ± 0.104, 4 | p = 0.460 |
| Interventricular Septum Thickness, Diastolic (mm) | 0.871 ± 0.047, 7 | 0.875 ± 0.048, 4 | p = 0.962 |
| Left Ventricular Mass | 119.6 ± 8.207, 7 | 156.3 ± 7.554, 4 | p = 0.016 |
| Fractional Shortening (%) | 41.54 ± 1.731, 7 | 26.95 ± 1.799, 4 | p = 0.0004 |
| Ejection Fraction (%) | 79.70 ± 1.791, 7 | 60.80 ± 2.801, 4 | p = 0.0002 |
| E wave Velocity (m/s) | 0.926 ± 0.017, 7 | 0.833 ± 0.034, 4 | p = 0.023 |
| Isovolumic Relaxation Time (ms) | 16.00 ± 0.873, 7 | 22.00 ± 2.582, 4 | p = 0.024 |
| Ejection Time (ms) | 56.00 ± 1.234, 7 | 54.00 ± 3.464, 4 | p = 0.524 |
| Isovolumic Contraction Time (ms) | 8.00 ± 1.234, 7 | 8.00 ± 0.00, 4 | p > 0.999 |
| Myocardial Performance Index | 0.429 ± 0.033, 7 | 0.563 ± 0.064, 4 | p = 0.0674 |

While changes in systolic function were most striking, diastolic function was also impaired but to a lesser extent. Mitral valve E-wave velocity was decreased 10% from 0.926 m/s in CON animals to 0.833 m/s in cKO mice (p=0.023; **Table** 4), suggesting diminution of peak inflow across the mitral valve in the early, passive phase of diastolic filling. Additionally, isovolumic relaxation time was prolonged from 16.00 to 22.00 ms in CON and cKO animals, respectively (p=0.024), without significant impact on ejection time or isovolumic contraction time (**Table 4**). While a consequent change in myocardial performance index failed to reach statistical significance, a trend toward an increase was observed (p=0.0674; **Table 4**), driven by the isovolumic relaxation time prolongation and suggesting reduced left ventricular diastolic function. These findings cumulatively point toward an impairment in cardiac function which commonly accompanies eccentric remodeling. This functional change was also predicted by proteomic pathway analysis, detecting diminution of proteins involved with cardiac muscle contraction (Figure 2C).

### Electrocardiographic changes in PFKFB2 cKO hearts

While we observe changes in metabolic profile and cardiac function, these were not necessarily sufficient to explain the early death phenotype of cKO mice. Electrocardiography was therefore performed, which revealed prolongation of parameters associated with ventricular conduction. Example tracings depict these genotype-dependent changes (Figure 6A). Specifically, durations of QRS and JT intervals, which represent depolarization and repolarization of the heart respectively, were increased (p=0.005-0.009; Figure 6B). This resulted in a 25.7% increase in QT interval (p=0.003; Figure 6B). As expected in the absence of a difference in RR interval between genotypes (Figure 6C), corrected QT interval was comparably increased (27.7%; p=0.003; Figure 6D). The observed QTc prolongation is likely attributable in large part to a temporal dispersion of repolarization. This is evidenced by a decrease in early repolarization, indicated by a 6.31-fold decrease in J wave amplitude (p<0.0001; Figure 6E), followed by a marked 74.6% increase in T_peak_-T_end_ interval (p=0.0047; Figure 6F), a strong predictor of sudden cardiac death [49]. Though, no difference was observed in T wave amplitude (Figure 6E). Taken together, these changes demonstrate a prolonged repolarization phenotype which is associated with predisposition to ventricular tachyarrhythmias, with QTc a well-established predictor of ventricular arrhythmia and all-cause mortality [20, 50, 51]. Ventricular specificity of these changes was suggested by unimpacted measures including PR interval and P wave duration (Figure 6G-H). These results support that electrophysiological derangements in the ventricular myocardium may underlie the sudden cardiac death phenotype observed in cKO animals.

**Figure 6.**
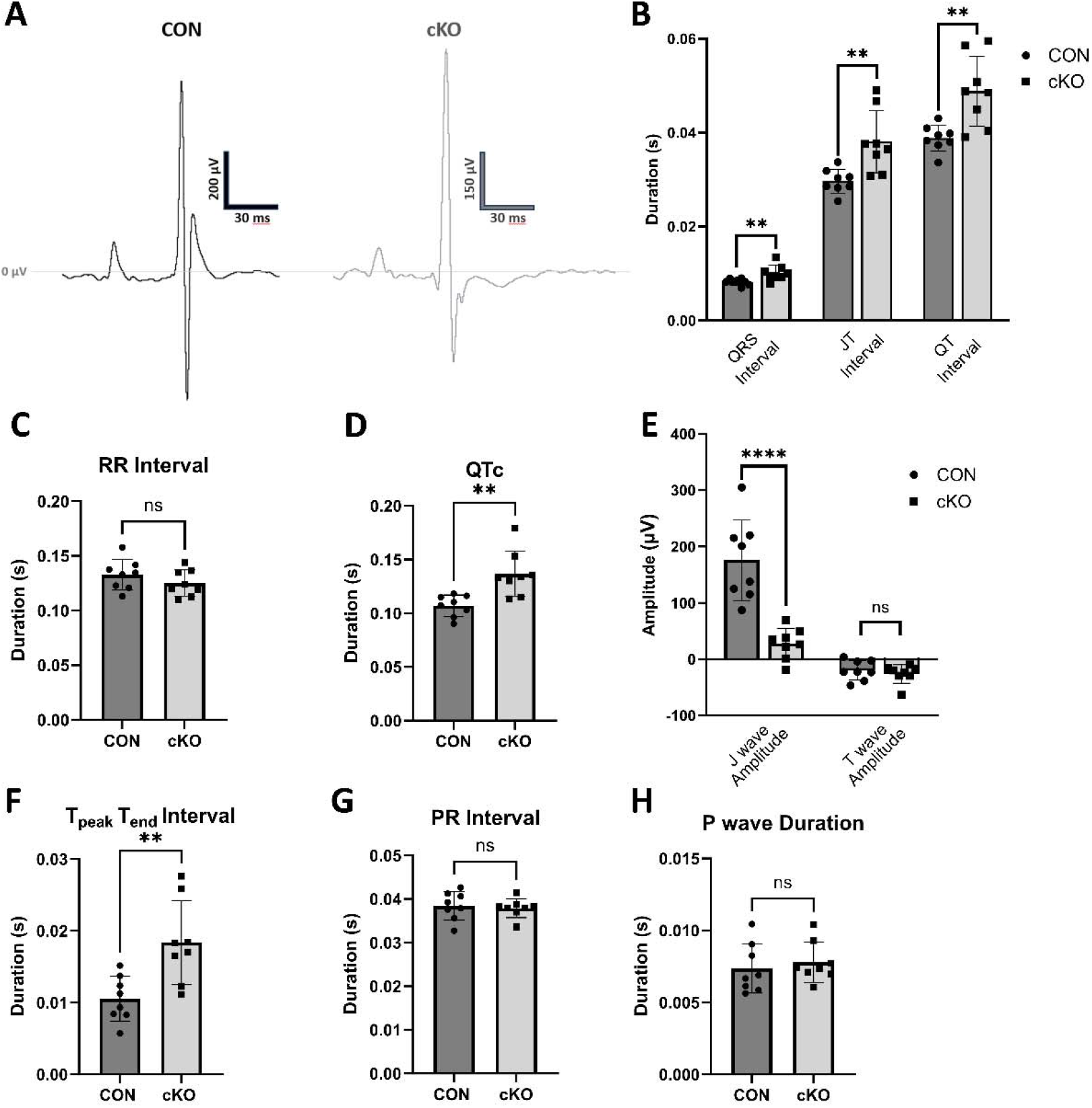
Ventricular conduction is affected at baseline in *Pfkfb2* knockout (cKO) animals relative to controls (CON). **A**, Representative tracing from cKO and CON animals. **B**, Interval durations associated with ventricular conduction (n=8/group). **C**, Average interval between R waves is unchanged between cKO and CON animals, indicating comparable heart rate (n=8/group). **D**, Corrected QT interval (n=8/group) calculated using Bazett Formula, QT_c_=QT/(RR Interval)^½^. **E**, Amplitudes of J and T waves (n=8/group). **F**, T_peak_T_end_ interval (n=8/group). **G**, PR interval (n=8/group). **H**, P wave duration (n=8/group). Data are shown as mean ± SD. Not significant (ns): p>0.05, **p≤0.01, ****p≤0.0001, unpaired Student’s *t* test.

## Discussion

This work demonstrates that loss of cardiac PFKFB2 drives metabolic reprogramming marked by changes suggestive of increased glucose oxidation, decreased fatty acid oxidation, and decreased mitochondrial abundance. These biochemical alterations were accompanied by pathologic cardiac structural and functional changes, with dilation and reduced systolic function among the most profound. Pathophysiological remodeling, as well as electrophysiological changes such as QT interval prolongation in cKO animals, are consistent with those often observed in heart failure of various etiologies.

Echocardiographic data from cKO animals demonstrates moderate dilation and an associated reduction in systolic function. Interestingly, previous work has shown that diminished levels of fructose 2,6-bisphosphate in an aortic-constriction-induced pressure overload model resulted in exacerbated cardiac remodeling and dysfunction [52]. This corroborates dilation and reduced function observed in cKO animals. Together, these data support the notion that normal glycolytic regulation by PFKFB2 is not only protective against pathology under pressure overload conditions but is also necessary for the preservation of normal cardiac structure and function under basal conditions. Further, metabolic changes observed in cKO animals strikingly mimic those observed in heart failure with reduced ejection fraction (HFrEF) and dilated cardiomyopathy. These changes are marked by a preferential shift from aerobic means such as fatty acid oxidation to carbohydrate catabolism and a decrease in amino acid metabolism [53, 54]. Interestingly, the left ventricular ejection fraction remained preserved in the cKO mice, raising the intriguing possibility that PFKFB2 knockout leads to a phenotype of electrical instability observed in HFrEF, even before severe structural changes occur. These changes in cKO animals and HFrEF are also interestingly accompanied by increases in metabolic kinases, such as AMPK, and include similarities such as increased GLUT1 abundance [53, 55, 56]. However, differences between these phenotypes do exist. Notably, while these other dilated cardiac pathologies are marked by increases in glycolysis and upstream carbohydrate catabolism, this does not always culminate in increased glucose oxidation [57, 58]. This uncoupling is thought to occur at the point of PDH activity which is not necessarily compensatorily upregulated in heart failure systems as it is in cKO animals [57]. Possible explanations for these differences in PDH activity are decreased PDK isoform expression and the increased insulin signaling and mitochondrial Akt association in cKO hearts.

There are pathophysiological similarities between cKO animals and changes which are observed in diabetes. This is despite distinctly different metabolic phenotypes with cKO hearts exhibiting increased Akt signaling, glucose oxidation, and PDH activity. While many individuals with diabetic cardiomyopathy present with concentric hypertrophic changes, another subset experiences this previously described eccentric remodeling with impairment of systolic function, especially in cases of HFrEF [19]. Interestingly, impacts on left ventricular structure at diastole, as well as changes to E wave velocity and ejection fraction, are comparable in cKO animals to those observed in a streptozotocin-induced model of type 1 diabetes (T1D) [59].

Baseline electrocardiographic changes are also remarkably reminiscent of those reported in diabetes, which are largely attributed to temporal dispersion of repolarization at a single cell level in diabetic models [59]. A coupling of glycolysis with intracellular Na^+^, K^+^, and Ca^2+^ transport in the cardiomyocyte has been previously observed. These relationships are largely based on impacts of glycolysis-derived ATP, including alterations to activity of ATP-sensitive K^+^ channels [60, 61], the Na^+^/K^+^ ATPase pump [62], and sarcoplasmic reticulum-related Ca^2+^ handling [63, 64]. These studies support that glycolysis-derived ATP is preferentially used for many electrical stability maintenance mechanisms [65]. However, as glycolysis-derived ATP production is likely preserved or increased in the cKO model, albeit via non-canonical mechanisms, the described relationships do not explain electrophysiological changes in this model. Little work has investigated the direct impacts of glycolytic regulation at pathophysiologically relevant points such as the PFKFB2/PFK-1 regulatory nexus on electrophysiological changes. Therefore, this work identifies glycolytic regulation as a potential contributor to crucial metabolic underpinnings of delayed cardiac conductance, which likely contribute to sudden cardiac death phenotypes in the hypoinsulinemic state that is caused by T1D.

Among possible shared contributors to these structural, functional, and electrophysiological changes is activation of the hexosamine biosynthesis pathway in cKO animals. While it is thought that acute activation of this pathway may serve a protective role in the context of myocardial ischemia, chronic activation has been shown to have deleterious implications, driving pathologic remodeling, reduced contractility, and altered Ca^2+^ and K^+^ handling [16, 47, 48, 66]. Structural and functional impacts are largely mediated by activation of transcriptional regulators such as nuclear factor of activated T-cells (NFAT), a prominent driver of pathologic remodeling, especially in response to pressure overload [67]. O-GlcNAcylation, the post translational modification which serves as an endpoint of the hexosamine biosynthesis pathway, has also been shown to activate the Ca^2+^/Calmodulin activated protein kinase II (CAMKII) [12]. CAMKII activity drives alterations in intracellular Ca^2+^ and K^+^ handling and is linked to action potential duration, QT interval duration, and ultimately arrhythmogenic predisposition [12, 17, 68]. As these mechanisms play a role in diabetes [12] and O-GlcNAcylation is increased in cKO hearts, it is an intriguing possibility that activation of CAMKII also underlies alterations to ventricular repolarization and QT interval duration observed in cKO animals. Additionally, recent literature demonstrates O-GlcNAcylation and activation of the cardiac ryanodine receptor (RyR2) directly, which could promote pro-arrhythmogenic sarcoplasmic reticulum Ca^2+^ leak through CAMKII-independent means [69].

Hexosamine biosynthesis pathway activation in the cKO model is likely due to the shunting of fructose 6-phosphate, the glycolytic intermediate directly upstream of PFK-1, when the PFKFB2/PFK-1 regulatory nexus is disrupted. Similarly, impaired insulin signaling and subsequent inactivation and degradation of PFKFB2 could be contributing to hexosamine biosynthesis pathway activation by this same mechanism in diabetes. It is therefore unsurprising that many of the pathological changes exhibited by cKO animals are also observed in some individuals with diabetic cardiomyopathy [70]. This supports the notion that PFK-2 may play a critical role in coupling glycolytic regulation to cardiac electrophysiology and function. The pentose phosphate pathway, which was also activated in cKO animals, generates NADPH and serves a predominantly protective role in cardiac disease, but is often downregulated in diabetic cardiomyopathy [58]. Enhanced PPP activity may support the NADPH-dependent antioxidants that are upregulated in cKO hearts. However, enhanced antioxidant capacity in cKO hearts is clearly insufficient to mitigate pathology. Interestingly, we have recently shown that the cardiac overexpression of a phosphatase-null PFK-2, which results in increased fructose-2,6-bisphosphate production and elevated cardiac glycolysis (Glyco^Hi^ mice), mitigates increases in the hexosamine biosynthesis and oxidative pentose phosphate pathways in the heart in animals exposed to a long-term high fat diet [18]. This further establishes a nodal role of PFK-2 in regulation of glucose ancillary pathways.

A hallmark of diabetic cardiomyopathy, metabolic syndrome, and heart failure is metabolic inflexibility. As a primary glycolytic regulator, it would be expected that a loss of PFKFB2 would drive metabolic inflexibility. We expected this to manifest as decreased glycolysis and glucose oxidation with an increase in FAO because the pro-glycolytic kinase activity of PFKFB2 dominates over its phosphatase activity [71]. However, in this regard, cKO mice are more reminiscent of late stage heart failure conditions where dependence on glucose increases. Nonetheless, while temporarily sustainable at baseline, the simultaneous decreases in other pathways may promote a metabolic inflexibility characterized by glucose dependence. Further, it is plausible that this would be exacerbated by an inability of the cKO heart to respond acutely to stress via direct glycolytic means in the absence of PFKFB2.

A limitation of this study is that we only addressed the baseline implications of PFKFB2 loss in these animals. To better understand the multifaceted regulatory role of PFKFB2, future studies should involve fasting and catecholamine-mediated stress. As well as demonstrating roles of both glycemic status and acute adrenergic stress, these future studies may provide insight into the role of cardiac PFKFB2 in predisposition of diabetic patients to arrhythmogenesis. This is particularly relevant as glycemic status is believed to contribute to ventricular arrhythmogenesis, particularly in cases of QT interval prolongation. However, a contribution of concomitant insulinemic status and subsequent PFKFB2 degradation is not routinely considered in clinical settings and in vivo models. A second limitation of this study is the lack of differentiation between sexes. As preliminary results did not demonstrate notable differences between sexes, it was not deemed necessary to analyze these data independently. Finally, since we did not use continuous telemetry in this study, we cannot confirm that the cause of sudden death in these animals was ventricular arrhythmias. Nonetheless, this is the most likely explanation given their QT interval prolongation.

This work demonstrates PFKFB2-dependent pathological implications of aberrant glycolytic regulation which are consistent with certain disease states such as diabetes. While it has previously been thought that the pathological impacts of PFK-2 degradation in diabetic cardiomyopathy were attributable to decreased glycolysis, these data suggest a concomitant role of glycolytic intermediate re-routing to ancillary pathways such as the hexosamine biosynthesis pathway that have the capability to subsequently promote pathology on chronic activation. The role of PFKFB2 and other isoforms has not been extensively studied in various other cardiomyopathy etiologies. Thus, the implications of this work could be even more far reaching than considered here and this demonstrates a potential therapeutic role of PFKFB2 restoration.

## Abbreviations

ACC: acetyl CoA carboxylase
Akt: protein kinase B
AMPK: adenosine monophosphate-activated kinase
BSA: bovine serum albumin
CAMKII: Ca^2+^/Calmodulin-activated protein kinase II
cKO: *Pfkfb2* cardiomyocyte-specific knockout
CON: control
DIA: data independent acquisition
EF: ejection fraction
FS: fractional shortening
FAO: fatty acid oxidation
GC-MS: gas chromatography mass spectrometry
GFAT: glucosamine: fructose-6-phosphate amidotransferase
GLUT-1: glucose transporter 1
GSK-3: glycogen synthase kinase
HBP: hexosamine biosynthesis pathway
HFrEF: heart failure with reduced ejection fraction
HIF-1: hypoxia inducible factor-1
LC-MS: liquid chromatography mass spectrometry
LDS: lithium dodecyl sulfate
MOPS: 3- (N-morpholino)propanesulfonic acid
MPC2: mitochondrial pyruvate carrier 2
NFAT: nuclear factor of activated T-cells
PC: palmitoyl carnitine
PDH: pyruvate dehydrogenase
PDK: pyruvate dehydrogenase kinase
PFK-2: Phosphofructo-2-kinase/fructose-2,6-bisphosphatase
PK: proteinase K
PPP: pentose phosphate pathway
Pyr: pyruvate
QT_c_: corrected QT interval
RT-qPCR: reverse transcriptase-quantitative polymerase chain reaction
SDS: sodium dodecyl sulfate
SRM: selected reaction monitoring
STZ: streptozotocin
TCA: tricarboxylic acid
TOM20: translocase of the outer mitochondrial membrane
T1D: type 1 diabetes

## Acknowledgements

We thank Melinda West and Yi Zhang for assistance with mouse colony establishment.

## Supplemental Material

Figures S1-13

(intended for publication as an online data supplement)

