## Supplementary material for "Loss of cardiac PFKFB2 drives Metabolic, Functional, and Electrophysiological Remodeling in the Heart": Supplmental Figures

**Figure S1**

**Few changes are observed in glucose metabolism or TCA cycle intermediates at the metabolite level, but nicotinamide metabolism is affected in response to *Pfkfb2* knockout.** Abundance of glucose, glycolytic intermediates, TCA and related intermediates, glucose derivatives, nicotinamide, and NAD^+^ were quantified using liquid chromatography (**B-I, L-M, O, Q-T**; n=5/group) and gas chromatography (**A, J-K, N, P, U-V**; n=5-6/group)-mass spectrometry in *Pfkfb2* knockout (cKO) and control (CON) animals. Relative abundance of metabolites was evaluated via unpaired Student’s *t* test without logarithmic transformation. Data are shown as mean ± SD. Not significant (ns): p>0.05, *p ≤ 0.05, **p≤0.01.

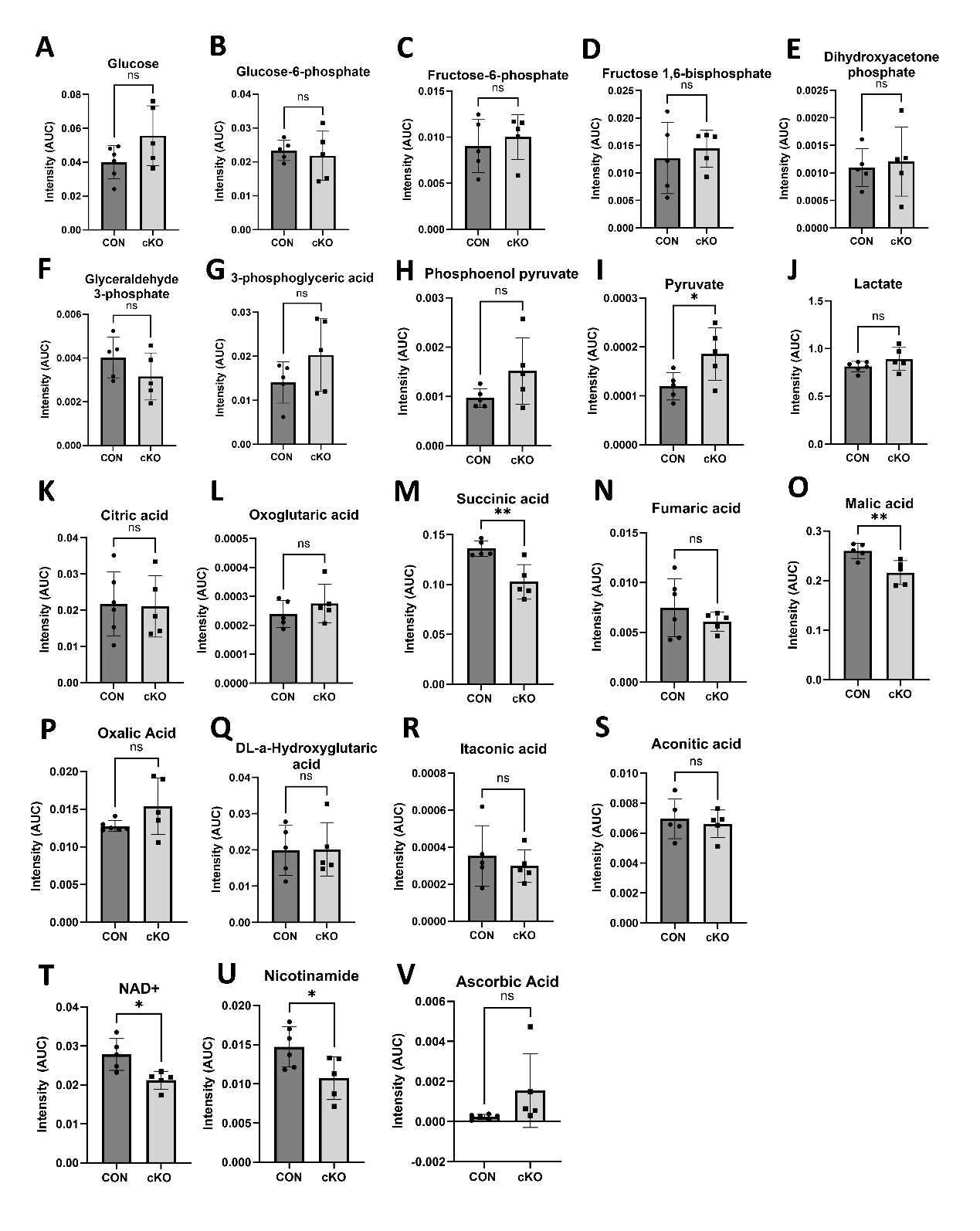

**Figure S2**

**Abundances of several amino acids are increased in response to *Pfkfb2* knockout.** Amino acid abundance was measured using liquid chromatography (**B-C, F, H, L-M, Q-R**; n=5/group) or gas chromatography (**A, D-E, G, I-K, N-P, S**; n=5-6/group)-mass spectrometry in *Pfkfb2* knockout (cKO) and control (CON) animals. Relative abundance of metabolites was evaluated via unpaired Student’s *t* test without logarithmic transformation (A-S). Data are shown as mean ± SD. Not significant (ns): p>0.05, *p ≤ 0.05, **p≤0.01, ***p≤0.001. Significantly different metabolites were subjected to pathway analysis and labeled pathways were found to be significantly affected (T).
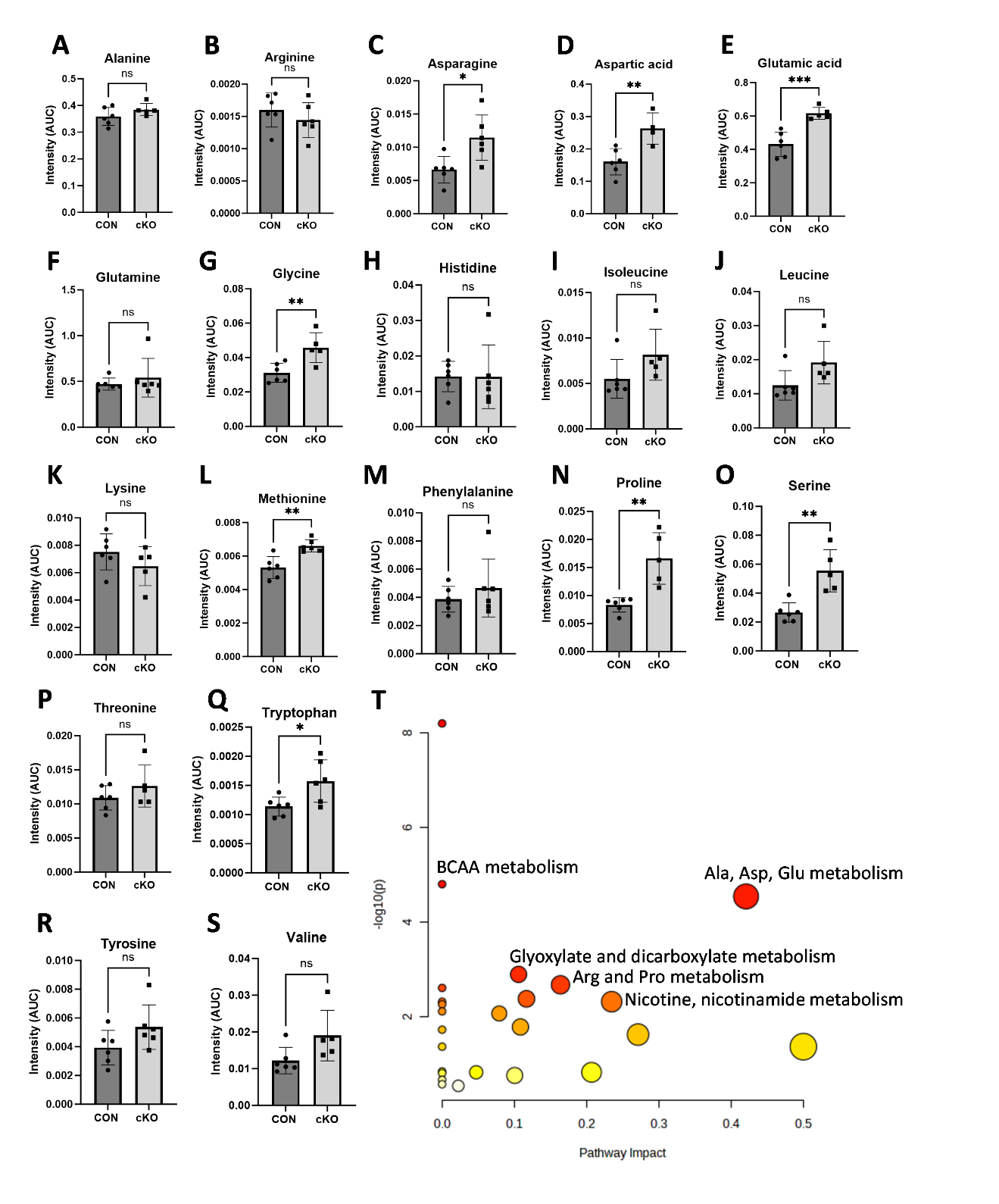

**Figure S3**

**Impact of *Pfkfb2* knockout on proteins involved with glucose metabolism.** Mass spectrometry was used to measure relative abundance of glucose transporters as well as glycolytic and related enzymes in *Pfkfb2* knockout (cKO; n=8) and control (CON; n=10) animals (**A-M**)**.** Data are shown as mean ± SD. Not significant (ns): p>0.05, *p ≤ 0.05, ***p≤0.001, ****p≤0.0001, unpaired Student’s *t* test.

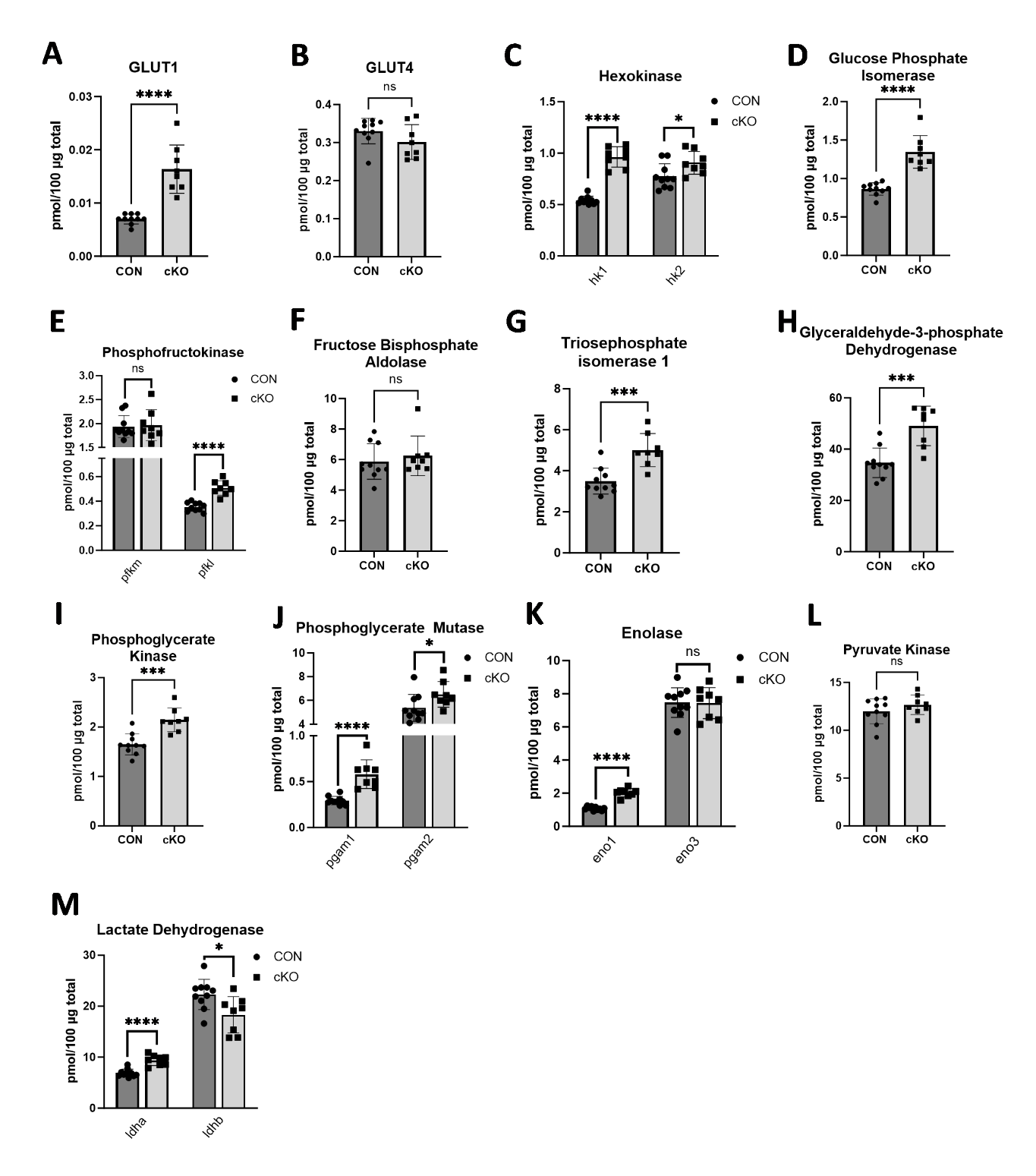

**Figure S4**

***Pfkfb2* knockout results in decreased TCA cycle enzyme abundance.** Mass spectrometry was used to measure relative abundance of TCA cycle enzymes in *Pfkfb2* knockout (cKO; n=8) and control (CON; n=10) animals. Data are shown as mean ± SD. Not significant (ns): p>0.05, *p ≤ 0.05, **p≤0.01, ***p≤0.001, ****p≤0.0001, unpaired Student’s *t* test.

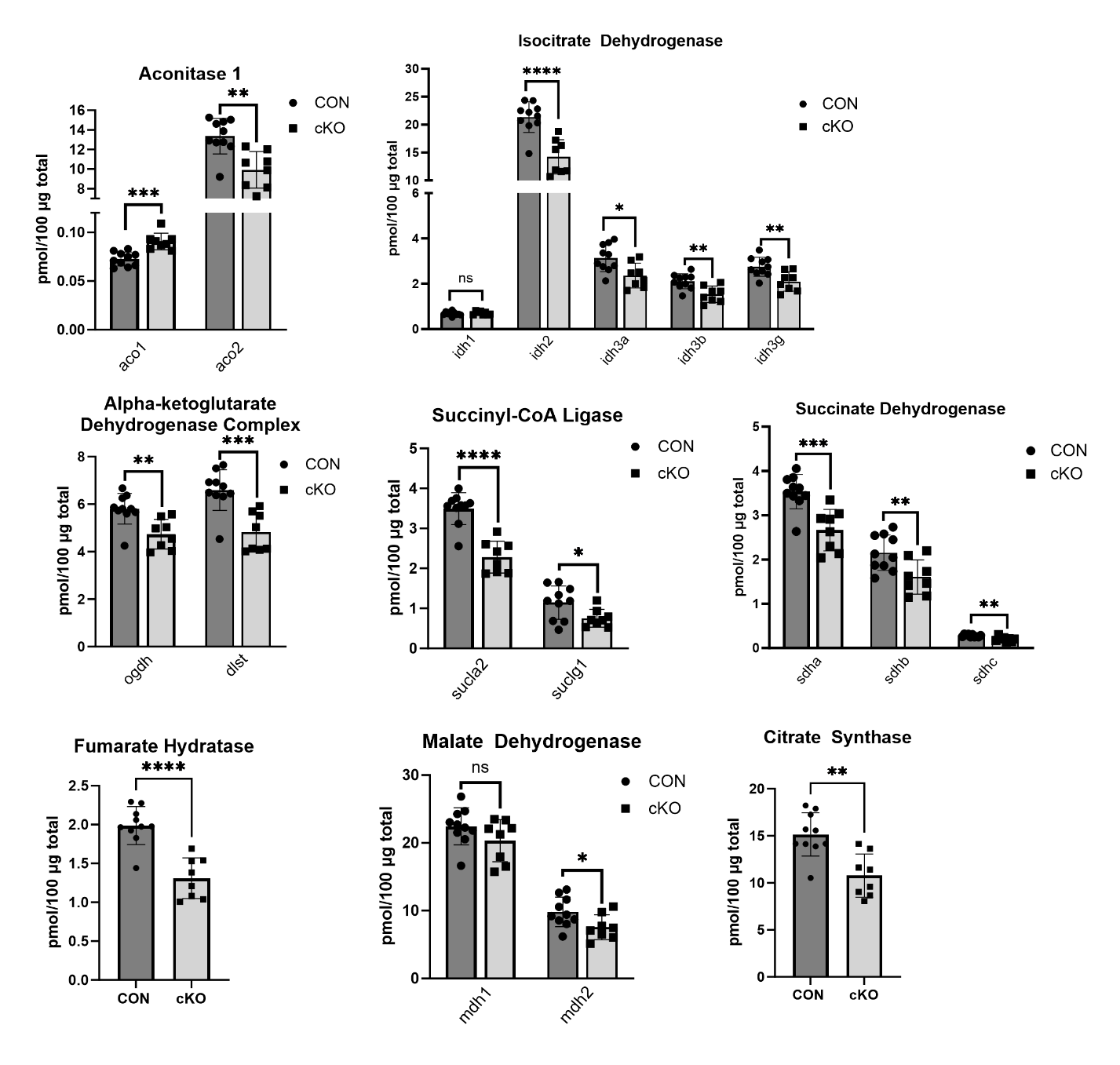

**Figure S5**

***Pfkfb2* knockout results in decreased mitochondrial complex I abundance.** Mass spectrometry was used to measure relative abundance of mitochondrial complex I subunits in *Pfkfb2* knockout (cKO; n=8) and control (CON; n=10) animals. Data are shown as mean ± SD. Not significant (ns): p>0.05, *p ≤ 0.05, **p≤0.01, ***p≤0.001, ****p≤0.0001, unpaired Student’s *t* test.

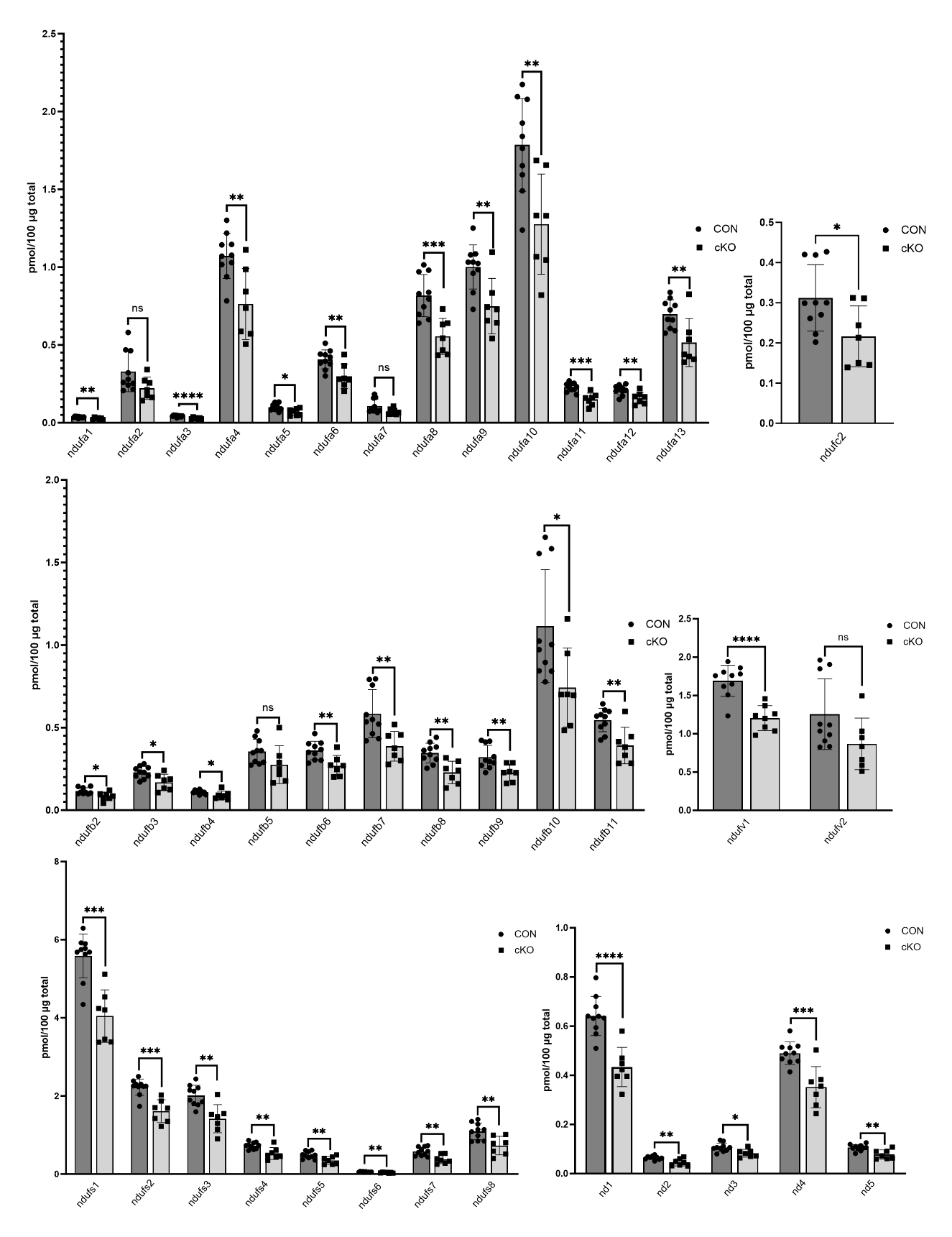

**Figure S6**

***Pfkfb2* knockout results in decreased abundance of electron transport chain components**. Mass spectrometry was used to measure relative abundance of mitochondrial complex III and IV subunits, ATP synthase subunits, cytochrome b, and cytochrome c in *Pfkfb2* knockout (cKO; n=8) animals and controls (CON; n=10). Data are shown as mean ± SD. Not significant (ns): p>0.05, *p ≤ 0.05, **p≤0.01, ***p≤0.001, ****p≤0.0001, unpaired Student’s *t* test.
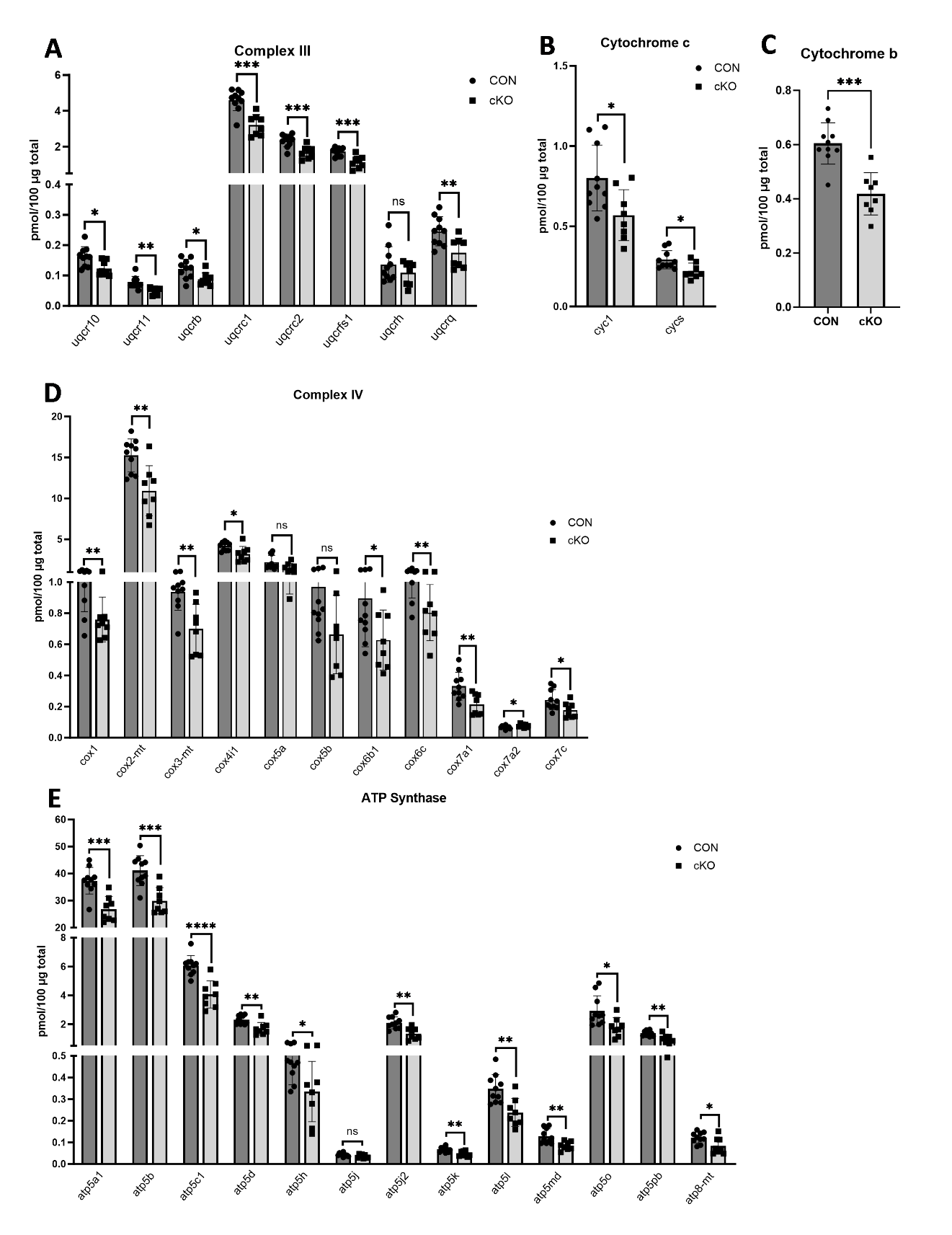

**Figure S7**

***Pfkfb2* knockout drives decreased abundance of proteins involved with beta oxidation.** Mass spectrometry was used to measure abundance of proteins involved with both mitochondrial (A-G) and peroxisomal (G-K) beta oxidation in *Pfkfb2* knockout (cKO; n=8) and control (CON; n=10) animals. Data are shown as mean ± SD. Not significant (ns): p>0.05, *p ≤ 0.05, **p≤0.01, ***p≤0.001, ****p≤0.0001, unpaired Student’s *t* test.

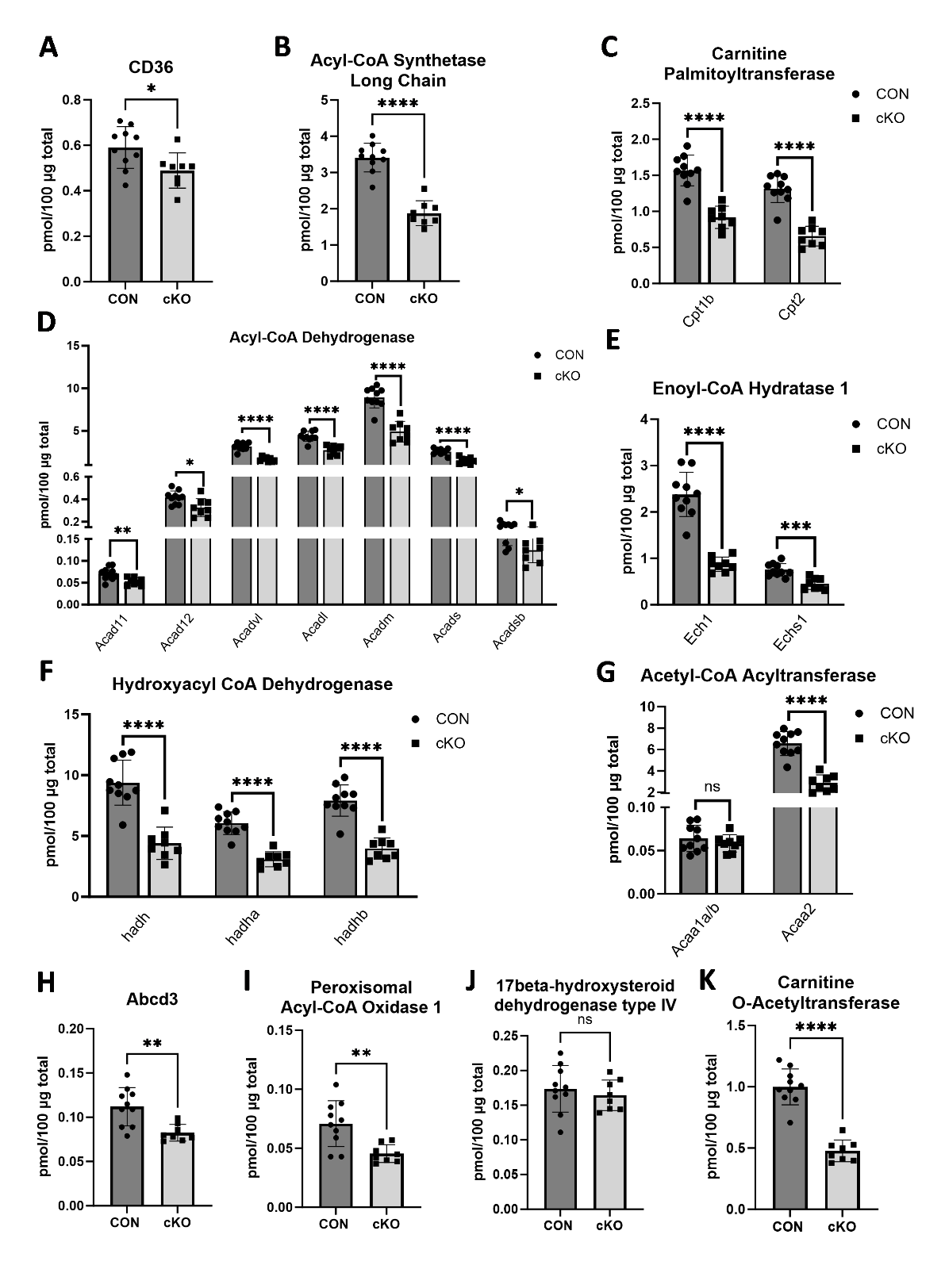

**Figure S8**

***Pfkfb2* knockout drives decreases in an array of proteins involved with fatty acid metabolism.** Mass spectrometry was used to measure abundance of proteins involved with fatty acid metabolism in *Pfkfb2* knockout (cKO; n=8) and control (CON; n=10) animals. Data are shown as mean ± SD. Not significant (ns): p>0.05, *p ≤ 0.05, **p≤0.01, ***p≤0.001, ****p≤0.0001, unpaired Student‘s *t* test.

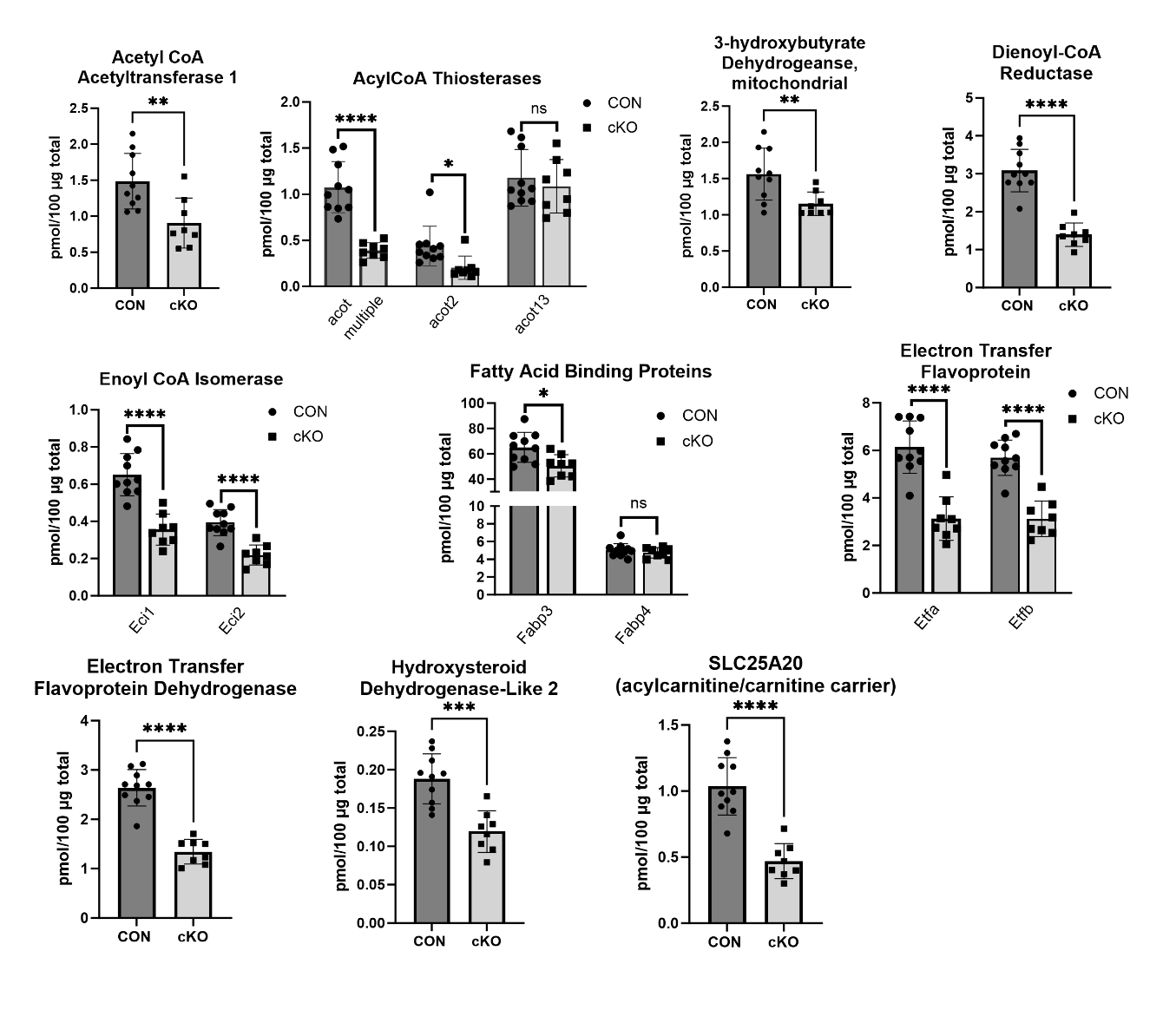

**Figure S9**

***Pfkfb2* knockout drives decreases in proteins involved with amino acid catabolism.** Mass spectrometry was used to measure abundance of proteins involved with amino acid catabolic reactions in *Pfkfb2* knockout (cKO; n=8) and control (CON; n=10) animals. Data are shown as mean ± SD. **p≤0.01, ***p≤0.001, ****p≤0.0001, unpaired Student’s *t* test.

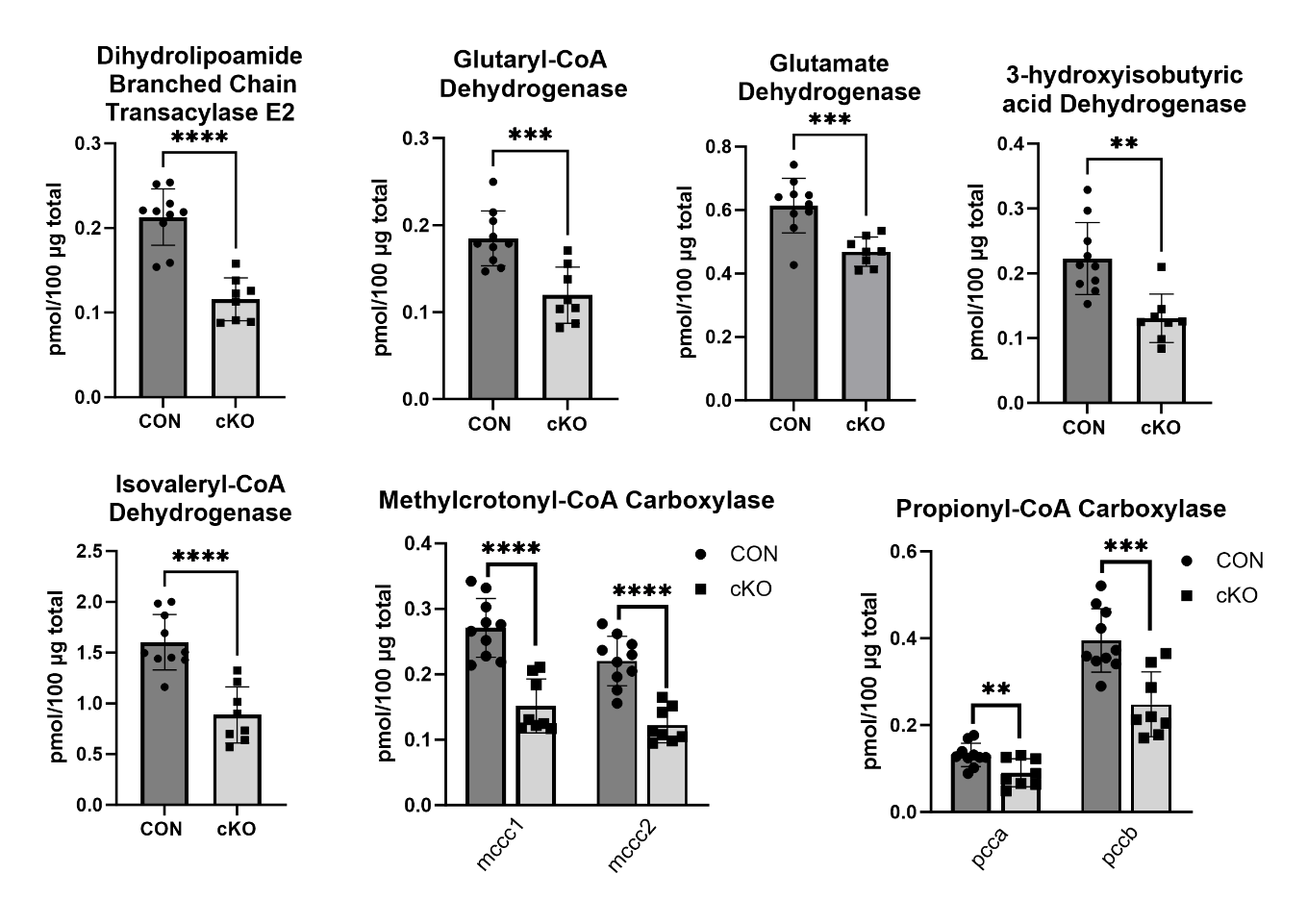

**Figure S10**

***Pfkfb2* knockout induces a broad increase in cellular responses to stress.** Mass spectrometry was used to measure abundance of proteins involved with antioxidant response (**A-G**) and chaperone proteins (**H-L**) in *Pfkfb2* knockout (cKO; n=8) and control (CON; n=10) animals. Data are shown as mean ± SD. Not significant (ns): p>0.05, *p ≤ 0.05, **p≤0.01, ***p≤0.001, ****p≤0.0001, unpaired Student’s *t* test.

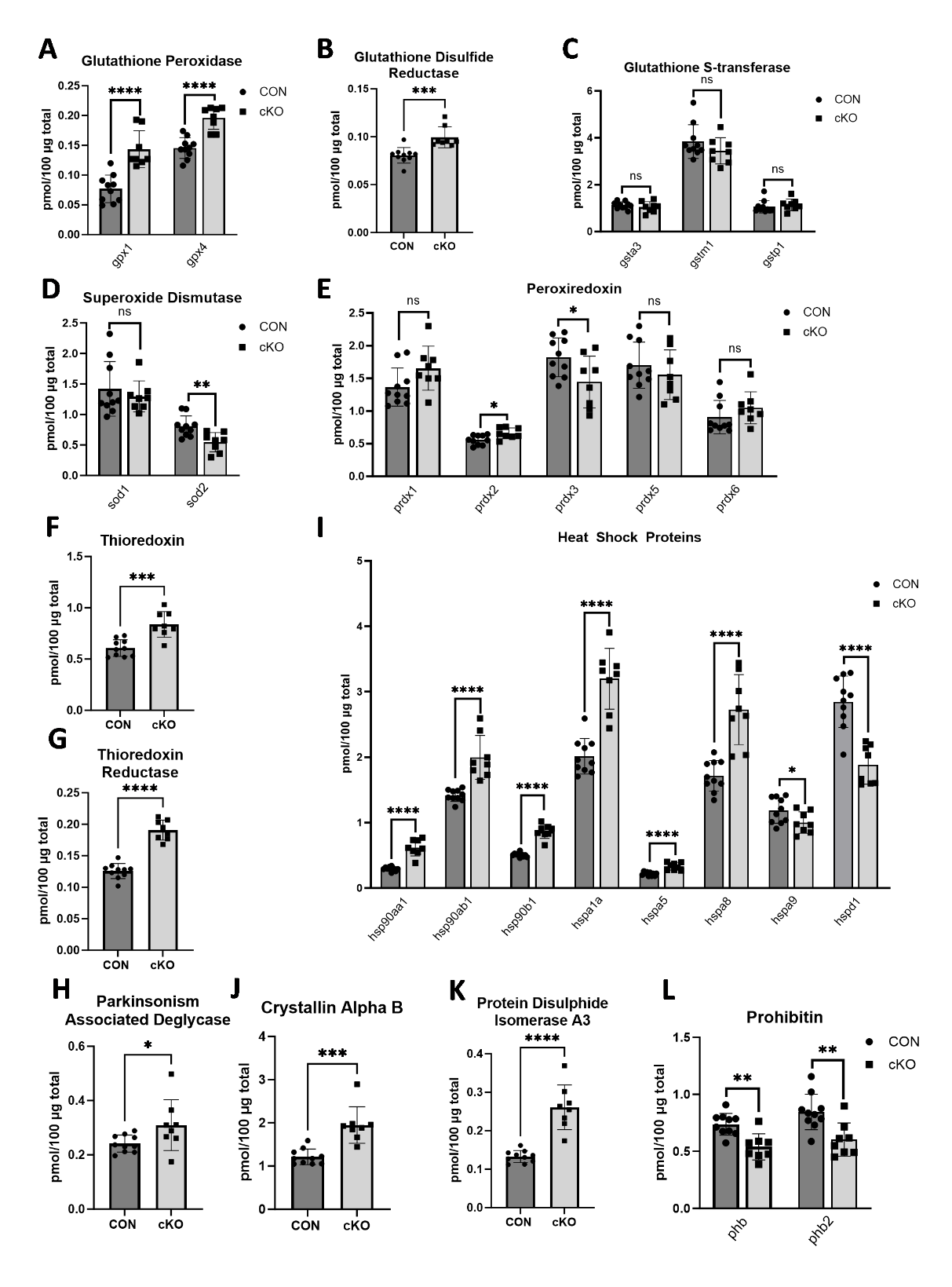

**Figure S11**

**Increased potential for protein turnover is observed in *Pfkfb2* knockout animals.** Mass spectrometry was used to measure abundance of eukaryotic translation elongation factor **(A)** and proteins involved with protein degradation (**B-F**) in *Pfkfb2* knockout (cKO; n=8) animals and controls (CON; n=10). Data are shown as mean ± SD. *p ≤ 0.05, **p≤0.01, ***p≤0.001, ****p≤0.0001, unpaired Student’s *t* test.

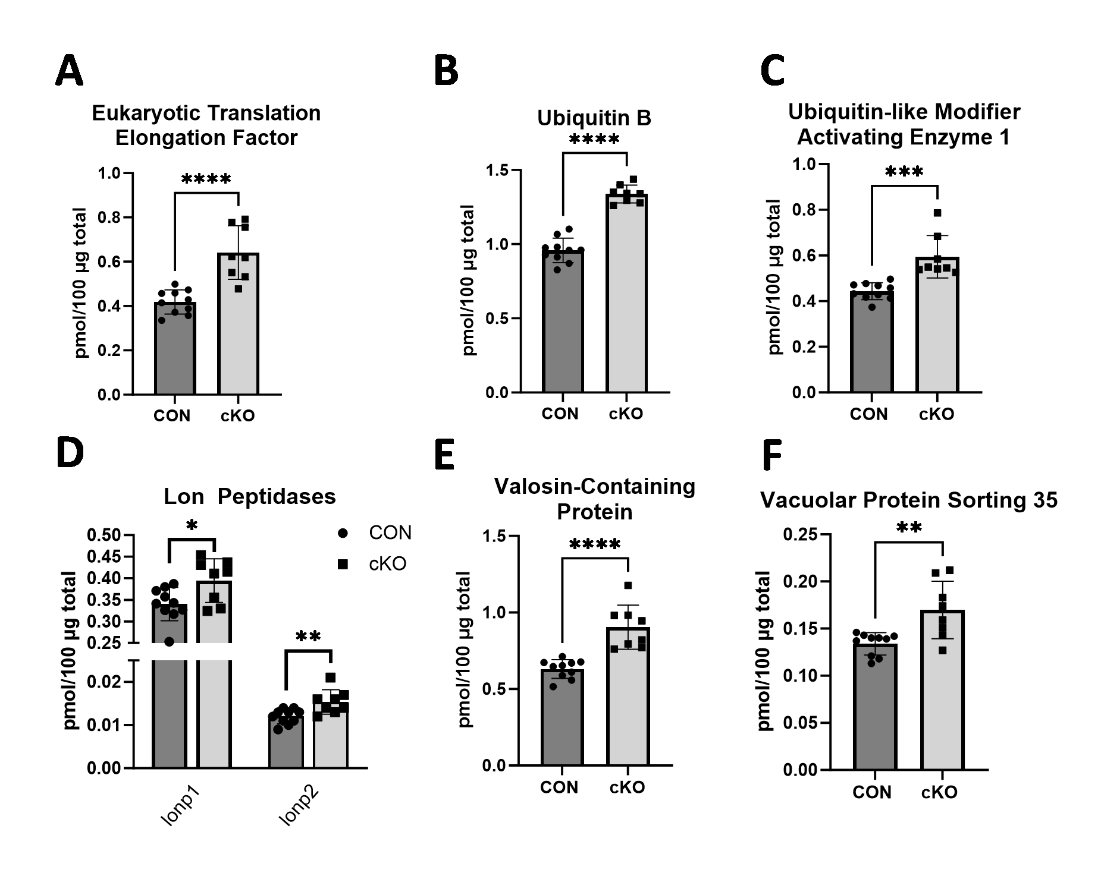

**Figure S12**

**Abundances of total and phosphorylated Acetyl CoA Carboxylase (ACC) are increased in *Pfkfb2* knockout animals.** Western blot analysis was used to measure abundances of ACC, and phosphorylated ACC (Ser79; p-ACC) and the ratio of p-ACC/ACC was calculated in *Pfkfb2* knockout (cKO; n=7) relative to control (CON; n=7) animals. Data are shown as mean ± SD. Not significant (ns): p>0.05, *p ≤ 0.05, unpaired Student’s *t* test.

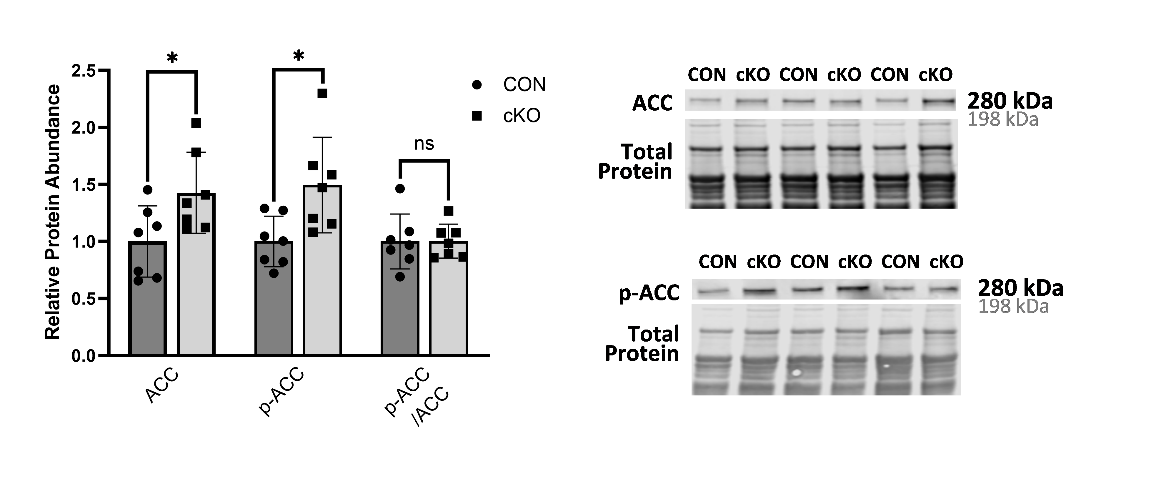

**Figure S13**

**Mitochondria-associated Akt is increased in *Pfkfb2* knockout animals.** **A,** Western blot analysis was used to measure abundance of Protein Kinase B (Akt) and phosphorylated Akt (Ser473; p-Akt) in the mitochondrial fractions of *Pfkfb2* knockout (cKO; n=7) relative to control (CON; n=7) animals and the ratio of p-Akt to Akt was calculated. Data are shown as mean ± SD. Not significant (ns): p>0.05, **p≤0.01, ***p≤0.001, unpaired Student’s *t* test. **B,** Mitochondria from *Pfkfb2* knockout (cKO; n=2) or control (CON; n=2) animals were treated with either proteinase K (PK), proteinase K and Triton X-100 (PK+Trition), or left un-treated. Western blot analysis was used to determine abundance of Pyruvate Dehydrogenase (PDH), Mitochondrial Pyruvate Carrier 2 (MPC2), Cytochrome c, Translocase of the Outer Mitochondrial Membrane (TOM20), Akt, and p-Akt. Abundance of these proteins in each treatment group was normalized to abundance in untreated mitochondria from that animal.

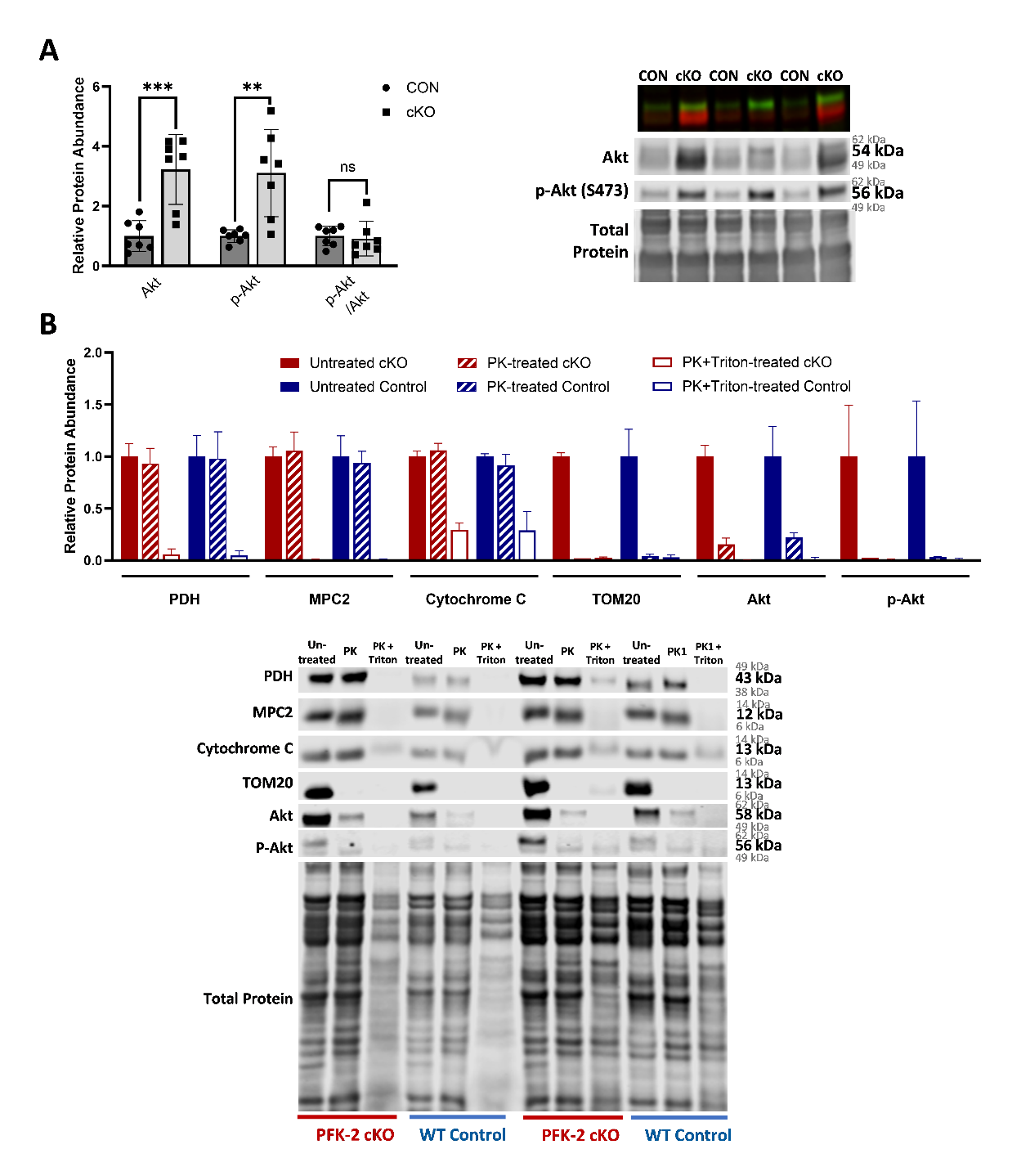
